# An ancient anthozoan protein reveals an alternative evolutionary path of antiviral signaling

**DOI:** 10.1101/2025.11.22.689925

**Authors:** Ton Sharoni, Adrian Jaimes-Becerra, Sydney Birch, Hee-Jin Kwak, Daria Aleshkina, Magda Lewandowska, Joachim M. Surm, Hannah Justin, Reuven Aharoni, Adam M. Reitzel, Yehu Moran

**Author notes:** Correspondence (YM), (TS).

## Abstract

How antiviral immunity first arose in animals is a central question in evolutionary biology. Using the sea anemone *Nematostella vectensis*, we identify *CARDIB*, a previously uncharacterized gene located next to *RLRb*, a cnidarian homolog of the vertebrate RIG-I-like receptor family. This conserved genomic linkage across Anthozoa reveals an ancient coupling between immune sensing and regulation. Despite sequence similarity to vertebrate MAVS, CARDIB performs an opposing function: it represses immune genes under basal conditions yet is essential for activation upon viral challenge. CARDIB binds RLRb via a single CARD domain, forming a repressive complex. Loss of either gene abolishes antiviral transcription, disrupts apoptosis, and elevates viral load under lab conditions. Both genes as well as the *RLRb* paralog, *RLRa*, are essential for antiviral defense under native conditions. Phylogeny places the cnidarian CARDs distinctly from the vertebrate RLR-MAVS families, revealing an ancient antiviral system regulating antiviral response through CARD-based signaling.

## Introduction

The innate immune system represents one of the most dynamic and rapidly evolving biological networks, continuously shaped by the tremendous evolutionary pressure of the arms race between hosts and viruses^1–3^. Identifying evolutionarily conserved antiviral defense mechanisms across distantly related animal lineages provides a critical framework for tracing the ancestral origins of immune recognition and signaling pathways. At the core of these systems are cytosolic RNA sensing receptors, which function as the primary detectors of viral RNA in vertebrates^4–6^. These receptors employ their C-terminal helicase domains to bind dsRNA and utilize the N-terminal caspase activation and recruitment domain (CARD) to engage downstream signaling partners. Two of these receptors, RIG-I and MDA5, sense and bind viral RNA that then exposes the CARDs to mediate direct interaction with a major component that is necessary for the activation of the immune response, the Mitochondrial Antiviral Signaling protein (MAVS). This protein is also known as IFN-β promoter stimulator I (IPS-1), caspase activation recruitment domain adaptor inducing IFN-β (CARDIF), or virus-induced signaling adaptor (VISA)^7^. The binding between the CARD domains of the RLRs and MAVS leads to MAVS polymerization into prion-like aggregates that amplify antiviral signaling^8^. This activation triggers the phosphorylation of the transcription factors Interferon Regulatory Factors 3/7 (IRF3/7) and the nuclear translocation of NF-κB, culminating in the induction of type I interferons (IFN) and proinflammatory cytokines^7,9^.

In mammals, the MAVS adaptor serves as a molecular hub in this signaling cascade. Located on the outer mitochondrial membrane, MAVS integrates the CARD-mediated input from activated RLRs and propagates the antiviral signal through self-assembly into filamentous aggregates^8,9^. Moreover, an increase in *MAVS* expression leads to a positive feedback loop that amplifies the immune response. On the other hand, decreasing *MAVS* leads to a reduction in the immune response^10–12^. Because of this central role, MAVS is a common target of viral antagonism. Many viruses express components that cleave or inhibit MAVS, thereby disrupting its mitochondrial localization, suppressing IFN production, and dampening the host’s antiviral signaling^13–16^. Thus, the RLR-MAVS signaling axis exemplifies a finely balanced molecular system in which structural cooperativity and evolutionary constraint coexist with persistent viral-driven diversification.

Despite extensive characterization in vertebrates, the evolutionary origin of the RLR-MAVS pathway remains elusive. Therefore, early-branching metazoans, particularly Cnidaria (corals, hydroids, jellyfish, and sea anemones), provide an evolutionary bridge to investigate these ancestral pathways, as this phylum is the sister group to Bilateria, which comprises the vast majority of extant animals^17,18^. The sea anemone *Nematostella vectensis* has become a powerful model organism with diverse genetic manipulation tools for studying innate immunity and antiviral evolution^19–21^. In *N. vectensis*, two paralogs of RLRs (*RLRa* and *RLRb*) have been identified and are suggested to contribute to antiviral defense. Both receptors are transcriptionally induced upon stimulation with the viral mimic polyinosinic:polycytidylic acid (poly(I:C)), a synthetic analogue of long dsRNA. However, when tested for their ability to recognize and bind viral RNA features, only RLRb showed direct binding affinity to long dsRNA. In contrast, exposure to short 5′ppp-dsRNA, a RIG-I-specific ligand, did not elicit a significant immune response from either RLR. These findings suggest a functional distinction between the two receptors. RLRb functions as the primary viral RNA sensor, exhibiting binding specificity for long dsRNA, resembling the ligand preference of MDA5 from vertebrates, whereas RLRa possibly plays a different role in immune activation^22^.

Complementing these findings, the *N. vectensis* genome encodes a remarkably large set of immune-related genes, many of which are homologous to vertebrate counterparts. These include the IFN pathway components, cGAS-STING signalling molecules, and apoptosis regulators, suggesting that key antiviral mechanisms were already present in the last common ancestor of cnidarians and bilaterians^22–24^. Consistent with this genomic complexity, experimental stimulation of *N. vectensis* with poly(I:C) induces a robust antiviral-like transcriptional response, markedly upregulating these immune-related genes^22^. Despite the robust antiviral response triggered by poly(I:C), a definitive MAVS ortholog has not been reported to date in cnidarian genomes.

In this study, we aim to distinguish between two evolutionary scenarios: whether a true MAVS ortholog exists in cnidarians, indicating that the RLR-MAVS signaling axis represents an ancient antiviral innovation predating the cnidarian-bilaterian split, or whether cnidarians instead evolved a parallel pathway that only partially mirrors the molecular architecture of bilaterian antiviral signaling while adapting to distinct selective pressures.

## Results

### Characterization of the CARDIB role as a repressor of the immune system

We identified a gene in the genome of *N. vectensis* that is a candidate homolog of *MAVS* in *H. sapiens*^22^. Based on a best reciprocal BLAST hit with the human MAVS protein, with a sequence identity of 33% and similarity of 50%. It is adjacent to the *N. vectensis RLRb* in the genome **(Fig. 1a)**. The main difference between the two proteins lies in their structure: MAVS in *H. sapiens* and other vertebrates comprises three domains: an N-terminal CARD, a C-terminal mitochondrial transmembrane domain, and a _∼_400 amino acid linker that provides flexibility for CARD filament assembly on the mitochondrial surface^7,25^. By contrast, the candidate MAVS homolog in *N. vectensis* is smaller and contains only a CARD domain. This prompted us to further investigate whether these genes share a conserved function.

**Fig. 1.**
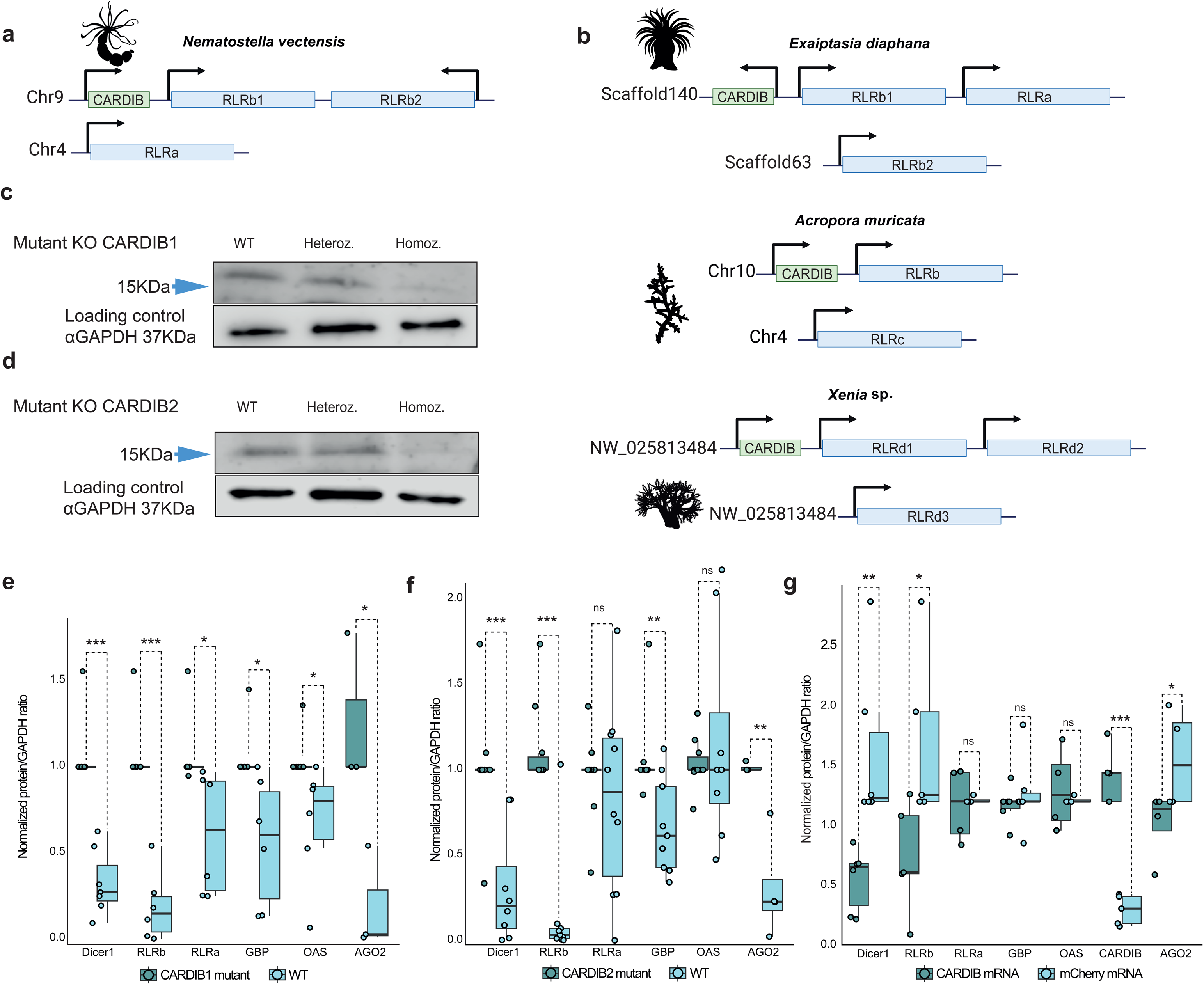
Conserved *RLRb-CARDIB* genomic organization in anthozoans, the generation of *CARDIB* mutant lines, and role in the immune response. **a,** Schematic representation of *RLRb-CARDIB* genomic organization in *N. vectensis.* b, Schematic structure of *RLRb-CARDIB* genomic organization in *E. diaphana*, *Xenia* sp., and *A. muricata*. Additional supporting information is available in **Supplementary file 3 and Supplementary** Fig. 4**. c-d,** Western blot validation of the loss of CARDIB protein, using an antibody against CARDIB tested on two CARDIB KO lines generated by the CRISPR/Cas9 approach based on two distinct non-overlapping guide RNAs (see methods). **e-f,** Western blots of different immune-related proteins (Dicer1, RLRb, RLRa, GBP, OAS, and AGO2) were measured in the mutant lines of CARDIB1 and CARDIB2 under basal conditions. Wild-type animals were used as a control. All the Western blot measurements were normalized to GAPDH protein, which served as a loading control. **g,** Microinjection of mRNA encoding CARDIB and mCherry separated by a self-cleaving P2A peptide into wild-type zygotes in order to overexpress CARDIB proteins. mCherry mRNA served as a control. The injected animals were analyzed 24 h post-injection. The same immune-related proteins mentioned above were tested by western blot, including CARDIB. All the Western blot measurements were normalized to GAPDH protein. Additional supporting information on antibodies against GBP, OAS, and CARDIB can be found in the **Supplementary file 1 Table 7**. Significance levels for **(e)**, **(f)**, and **(g)** were tested by paired two-tailed t-test; *p-value < 0.05, **p-value < 0.01, ***p-value < 0.001; NS, not significant **(Supplementary file 1 Tables 13-15)**.

For this, we next examined whether the *RLRb-MAVS*-like genomic organization, where the two genes are close neighbors, is conserved beyond *N. vectensis*. Our analysis revealed that this organization is present in representatives of all major anthozoan lineages, all the way to soft corals, indicating strong conservation of this genomic arrangement and possible shared transcriptional regulation of these neighboring genes **(Fig. 1b)**.

To characterize the function of this gene in *N. vectensis*, we generated two independent knockout lines using CRISPR/Cas9 technology, each targeting a different site within the gene. A successful knockout was verified at the protein level by Western blot **(Fig. 1c-d)**.

We first assessed the expression of immune-related homologous genes under basal conditions in adult mutants compared with wild-type animals. Unexpectedly, the absence of the MAVS-like gene led to the upregulation of multiple immune-related homologous genes, rather than a decrease in expression, like in mammals^9^ **(Fig. 1e-f)**.

To further investigate the function of this gene, we overexpressed the MAVS-like protein by injecting *N. vectensis* embryos with mRNA encoding MAVS-like and mCherry^26^ separated by a self-cleaving P2A peptide^27^, while mCherry-encoding mRNA served as a control. Complementing the knockout results, overexpression of MAVS-like led to a reduction in the protein levels of RLRb, DICER1, and AGO2 **(Fig. 1g)**, which are suspected to be central players in the initiation of antiviral immunity in *N. vectensis*^22^.

Taken together, these findings suggest that although this gene shows sequence similarity to MAVS, it exerts an opposite effect in *N. vectensis*, acting as a repressor of immune-related genes. Based on this role and on the further findings described below, we named the gene *CARD Inhibitor Binding protein* (*CARDIB*).

### CARDIB binds RLRb

Previous studies have shown that interactions between the CARD domains of MAVS and RLRs are essential for initiating the immune response in mammals^8,9^. In contrast, *CARDIB* in *N. vectensis* functions as a repressor of immune activity, although it still retains the structural features of a CARD domain. In mammals, the two major RLRs, RIG-I and MDA5, each carry two N-terminal CARD domains. Intriguingly, the *N. vectensis* homologs RLRa and RLRb are predicted by hidden-Markov model/domain annotations to have only a single CARD domain, lacking a detectable second CARD^22^. Yet, in the nematode *C. elegans*, despite limited primary-sequence similarity to canonical CARDs, the RLR homologs DRH-1 and DRH-3 have N-terminal regions that adopt tandem CARD folds, as was shown by the crystal structure of DRH-3^28^ and structure/function analysis of DRH-1^29^. Hence, to verify the apparent absence of a second CARD domain in RLRa and RLRb, we predicted their structures with AlphaFold2 **(Fig. 2a-b)**, which produced high-confidence superpositions to CARD1 of MDA5 but not to CARD2 (RMSD ≈1 Å vs >12 Å), supporting the absence of a second CARD-like fold in both sea anemone proteins.

**Fig. 2.**
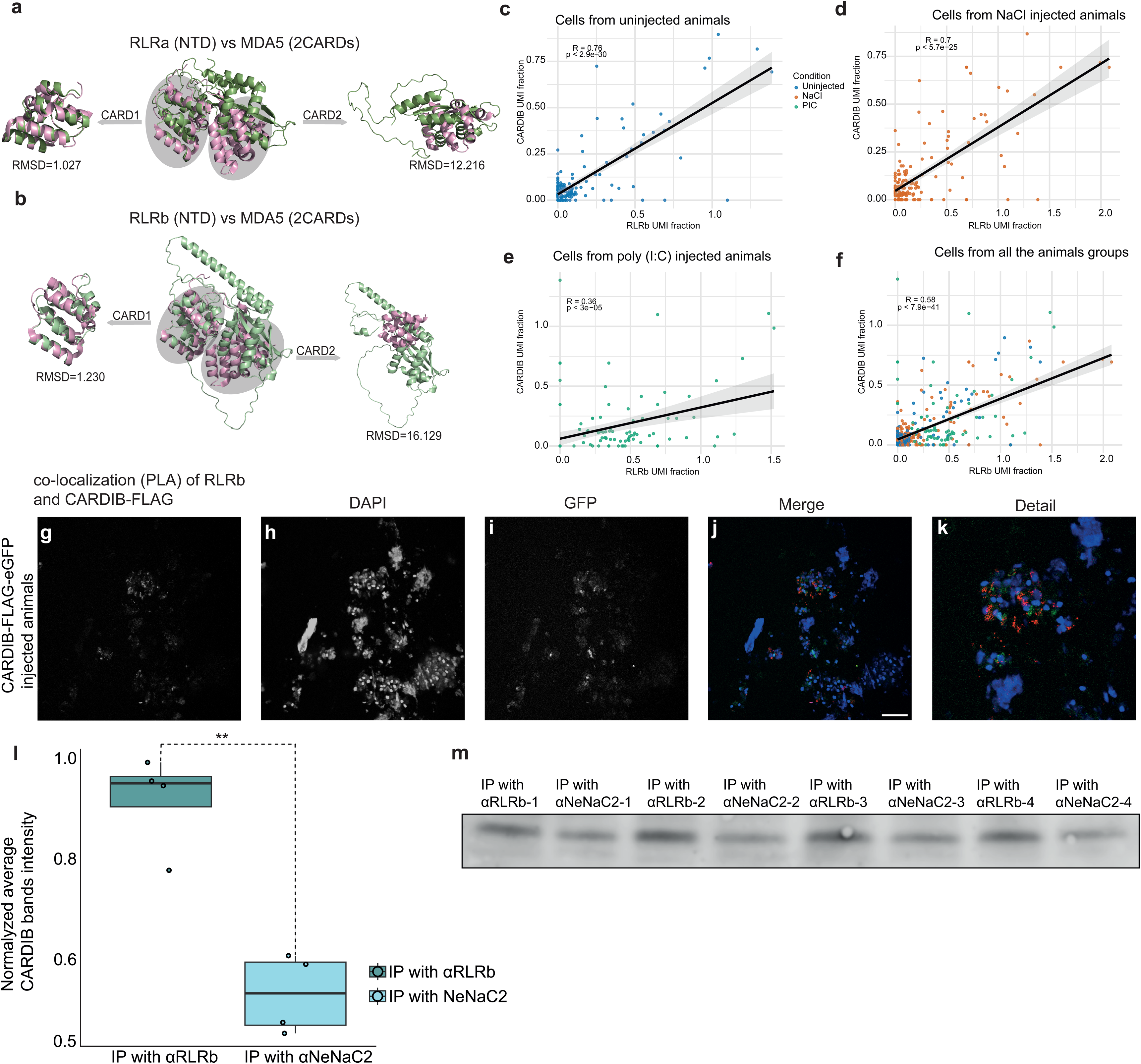
Interactions between CARDIB and RLRb, and the role of CARDIB in repressing the immune response. a-b, Predicted AlphaFold2 structures of *N. vectensis* RLRa and RLRb were superimposed with the MDA5 CARD domains, and structural similarity was evaluated by calculating the root mean square deviation (RMSD). **c-f,** Single-cell RNA sequencing (scRNA-seq) correlation analysis of the appearance of *CARDIB* and *RLRb* in the same cell in each of the different injection treatments, poly(I:C), NaCl, and uninjected^30^. **g-k,** Confocal images of proximity ligation assay (PLA) signals in dissociated cells derived from 24-h-old planulae injected with C-terminal CARDIB-FLAG-GFP mRNA; PLA signal for RLRb and FLAG co-localization (red); DAPI (blue); GFP (green); scale bars are 20 µm. Representative single z-stack section images were obtained from at least three independent replicates. Supporting information can be found in **Supplementary** Fig. 1. **l-m,** Co-immunoprecipitation of CARDIB and RLRb measured by western blot **(m)**. Band intensities were quantified with software (Li-COR Biosciences) and calculated as fold changes relative to the loading control for each sample **(l)**. Significance level was tested by paired two-tailed t-test; *p-value < 0.05, **p-value < 0.01, ***p-value < 0.001; NS, not significant. Additional supporting information is provided in **Supplementary file 1 Tables 16-17**.

Next, we examined the correlation between *CARDIB* and *RLRb* expression in the context of immune activation, using single-cell RNA-seq data from an experiment, in which *N. vectensis* transgenic reporter line expressing mCherry under the *RLRb* promoter was injected with poly(I:C) to activate the immune response; NaCl injected and uninjected animals served as controls^30^. Cells were analyzed 24 h post-injection. Correlation analysis showed that *CARDIB* and *RLRb* expression was stronger in uninjected, and NaCl-injected animals compared with poly(I:C) injected animals **(Fig. 2c-f)**. Thus, reduced correlation between *CARDIB* and *RLRb* expression was associated with stronger immune activation, consistent with our earlier findings that CARDIB represses multiple immune-related genes in *N. vectensis*.

To further test this interaction, we performed a proximity ligation assay (PLA) with antibodies against RLRb and FLAG tag in 24 h planulae injected with C-terminal CARDIB-FLAG-GFP mRNA at the zygote stage; GFP mRNA alone served as a control **(Supplementary Fig. 1)**. We observed multiple events of co-localization of the injected CARDIB-FLAG protein construct and endogenous RLRb protein. PLA results indicated a strong likelihood of interaction between CARDIB and RLRb **(Fig. 2g-k)**.

Finally, we examined CARDIB-RLRb interactions without overexpression. Total protein was extracted from wild-type 24-h-old embryos, and co-immunoprecipitation (Co-IP) was performed on the extracts using anti-RLRb antibodies, with anti-NeNaC2 antibodies serving as a control. Western blotting with anti-CARDIB antibodies showed a significant enrichment of CARDIB in samples immunoprecipitated with RLRb **(Fig. 2l-m)**.

Altogether, these results provide strong evidence that CARDIB directly interacts with RLRb, probably via the single CARD of this dsRNA receptor, supporting its role as a repressor of the immune response in *N. vectensis*.

### Clustering and phylogenetic analysis of CARD domains

To elucidate how cnidarian CARD domains relate to the broader metazoan CARD protein family, we combined clustering and phylogenetic approaches using 1,055 sequences representing six major animal lineages. This framework enabled us to evaluate whether *N. vectensis* CARD-containing proteins share ancestry with vertebrate immune receptors such as RIG-I, MDA5, and MAVS, or instead constitute a distinct lineage. A maximum-likelihood phylogeny reconstructed from these 1,055 CARD domain sequences revealed five well-supported clades corresponding to the CARD-containing proteins **(Fig. 3a)**. Node support was assessed using SH-aLRT, aBayes, and ultrafast bootstrap tests, shown as three values at each node. Most major clades exhibited high support across all three tests, confirming their robustness. In addition to these five clades, an additional cluster composed predominantly of sequences from *N. vectensis* (highlighted in orange) and other anthozoans were recovered with moderate support. Although the CLANS similarity network, based on pairwise BLAST similarities, also identified five principal clusters corresponding to the RIG-I, MDA5, and MAVS receptors **(Fig. 3b)**, it placed the cnidarian CARD sequences within the vertebrate MAVS group. In contrast, the phylogenetic reconstruction positioned these sequences as a separate lineage. Together, these results suggest that while cnidarian CARD domains retain sequence similarity to vertebrate MAVS, their evolutionary history likely reflects an independent diversification event or convergence rather than direct orthology.

**Fig. 3.**
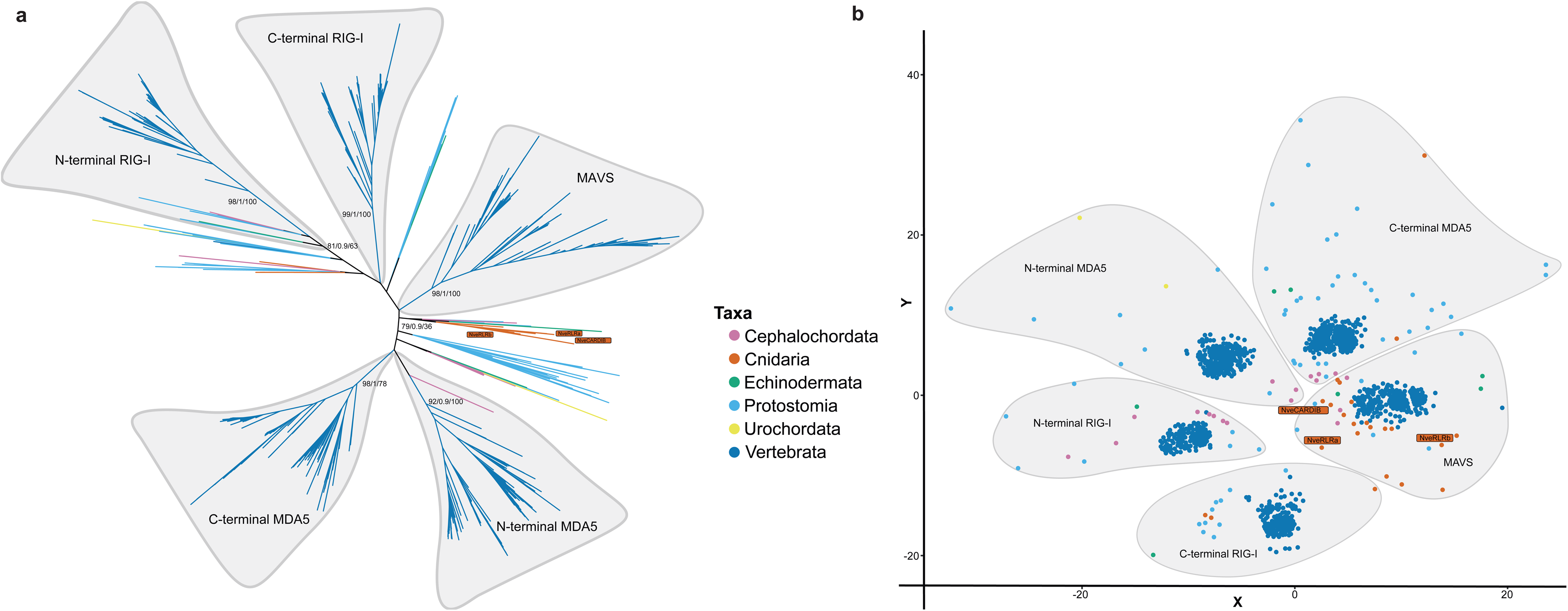
Phylogeny and clustering of CARDs from Anthozoa and other metazoans. **a,** Maximum-likelihood phylogeny of 1,055 CARD domain sequences from six metazoan groups: Cephalochordata, Cnidaria, Echinodermata, Protostomia, Urochordata, and Vertebrata. Values on nodes represent SH-aLRT/aBayes/bootstrap supports. The five main clades correspond to known CARD-containing proteins (RIG-I N-terminal, RIG-I C-terminal, MDA5 N-terminal, MDA5 C-terminal, and MAVS). An additional group containing *N. vectensis* sequences highlighted in orange (NveCARDIB, NveRLRa, and NveRLRb) forms a separate lineage. **b,** CLANS similarity network of the CARD dataset. Each point represents a CARD sequence color-coded by taxonomic group. The five principal clusters correspond to the N- and C-terminal CARDs of RIG-I and MDA5, as well as MAVS. *N. vectensis* sequences, highlighted in orange, are grouped within the MAVS cluster.

To further investigate why the *N. vectensis* CARDIB clade exhibits strong sequence similarity to vertebrate MAVS in the clustering network yet forms a closely related but distinct lineage in the phylogeny, we examined whether differences in selective pressures could account for these patterns. The branch model **(Supplementary file 1 Table 1)** showed no significant difference in selective pressure between foreground (MAVS clade and the *N. vectensis* CARDIB clade) and background lineages (the four other well-supported clades) (lnL = −19103.159 vs −19102.615; LRT = 1.09, p > 0.05). However, the estimated ω values indicated that the CARDIB and MAVS branches (ω = 0.10) evolved under stronger purifying selection than the remaining CARD-domain clades (ω = 0.26), suggesting slower overall evolutionary rates. In contrast, the branch-site model **(Supplementary file 1 Table 2)** detected significant episodic positive selection on the same foreground branches (lnL_alternative = −19036.418 vs lnL_null = −19040.735; LRT = 8.63, p < 0.01), with roughly 15% of sites (classes 2a + 2b) showing a very high ω_₂_ ≈ 227, while most codons remained highly constrained (ω_₀_ ≈ 0.26). This suggests that despite the overall sequence conservation due to purifying selection, some sites still evolve under strong positive selection, possibly due to being involved in an arms race with viral pathogens.

### Poly (I:C) microinjection to N. vectensis mutant vs wild-type

The discovery of *CARDIB* as a gene distinct from the mammalian *MAVS*, with an opposite role as a repressor of homologous immune-related genes, together with its ability to bind RLRb, led us to investigate the contribution of CARDIB to the immune response in *N. vectensis*. To this end, in addition to the two *CARDIB* knockout mutant lines we generated two knockout lines for *RLRa*, and one for *RLRb*. For the *RLRb* knockout line, both gene copies were disrupted simultaneously.

After generating the mutant lines, we verified by western blot the absence of the respective proteins in each mutant compared with wild-type animals **(Fig. 1c, d)** and **(Fig. 4a, b**). Zygotes of both homozygous mutants and wild-type animals were then injected with poly(I:C) to stimulate an antiviral response or with NaCl as a control. Transcriptome profiling was performed 24 h post-injection **(Fig. 4c-f)**. The analysis revealed that *CARDIB* and *RLRb* mutants showed almost no response to poly(I:C) compared with wild-type animals, whereas RLRa mutants displayed a nearly intact transcriptomic response similar to wild-type **(Fig. 4c-e)**.

**Fig. 4.**
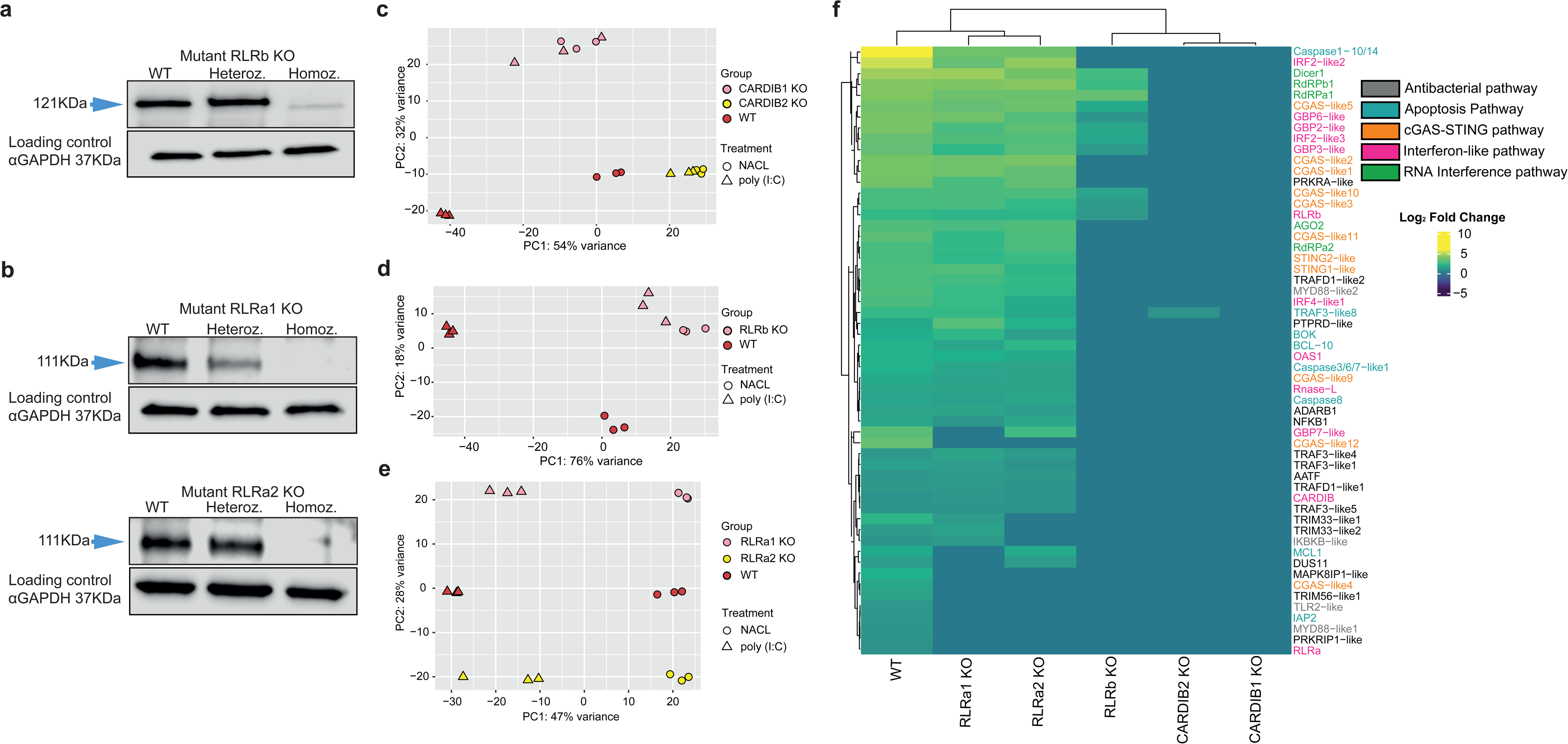
Knockout of *CARDIB* and *RLRb* suppresses the immune system of *N. vectensis*. a-b, Western blot validation shows the absence of each protein, RLRb, or RLRa, in their respective mutant lines. **c-e,** PCA plots illustrating the global transcriptomic profiles of poly(I:C) injected wild-type and mutant animals at 24 h post-injection. NaCl injected the animals served as controls. Three biological replicates were tested for each sample. **f,** A set of 56 immune-related genes was selected based on their previously reported upregulation following poly(I:C) injection in *N. vectensis* zygotes^22^. These genes were examined as indicators of immune activation and response efficiency of the immune response in the mutant animals of *CARDIB*, *RLRb*, *RLRa*, and wild-type after 24 h from the injection of poly(I:C) to their zygotes. The heatmap displays log_₂_ fold changes in gene expression in response to stress. The color scale ranges from −10 (strong downregulation) to +10 (strong upregulation): yellow indicates strong induction (log_₂_FC > 5), green reflects mild induction (log_₂_FC ≈ 1), and purple indicates downregulation (log_₂_FC < −5). Additional supporting information is provided in **Supplementary file 1 Table 18**

We next examined a previously curated set of homologous immune-related genes in *N. vectensis* that respond to poly(I:C) exposure and increased viral load in wild-type animals^31^ **(Fig. 4f).** In *CARDIB* and *RLRb* mutants, no significant differences were observed between animals injected with poly(I:C) and those injected with NaCl, confirming the essential role of these genes in mounting an antiviral response. In contrast, *RLRa* mutants exhibited expression patterns similar to wild-type animals, although the overall immune response appeared reduced.

Together, these results identify CARDIB and RLRb as major components required for the initiation of the antiviral immune response in *N. vectensis*. By comparison, the removal of RLRa somewhat diminished the strength of the response but did not abolish it.

### Viral load in CARDIB and RLR mutants

After revealing the importance of CARDIB and RLRb for initiating the immune response in *N. vectensis*, we next assessed how the functionality of the mutant immune system was disrupted by removing these key components. Specifically, we examined changes in viral load and apoptosis, a defense mechanism against intracellular pathogens triggered by immune challenge^23,32^ .

First, adult animals from the mutant and wild-type lines were grown for two weeks under identical conditions. Ribo-depleted libraries were generated to quantify viral load **(Fig. 5a-b)**. We first examined changes in the six most abundant viruses present in laboratory cultures of *N. vectensis*^22,31,33^, comparing each mutant line with wild-type animals **(Fig. 5a)**. *CARDIB* and *RLRb* mutants showed significantly higher susceptibility to most of these viruses relative to wild-type animals. By contrast, *RLRa* mutants displayed viral levels similar to those of wild-type animals. Along with measurement of core virus reads, we then examined the total viral load **(Fig. 5b)**. Consistent with the previous findings, *CARDIB* and *RLRb* mutants exhibited a significant increase in total viral load compared with wild-type, whereas *RLRa* mutants showed no significant change.

**Fig. 5.**
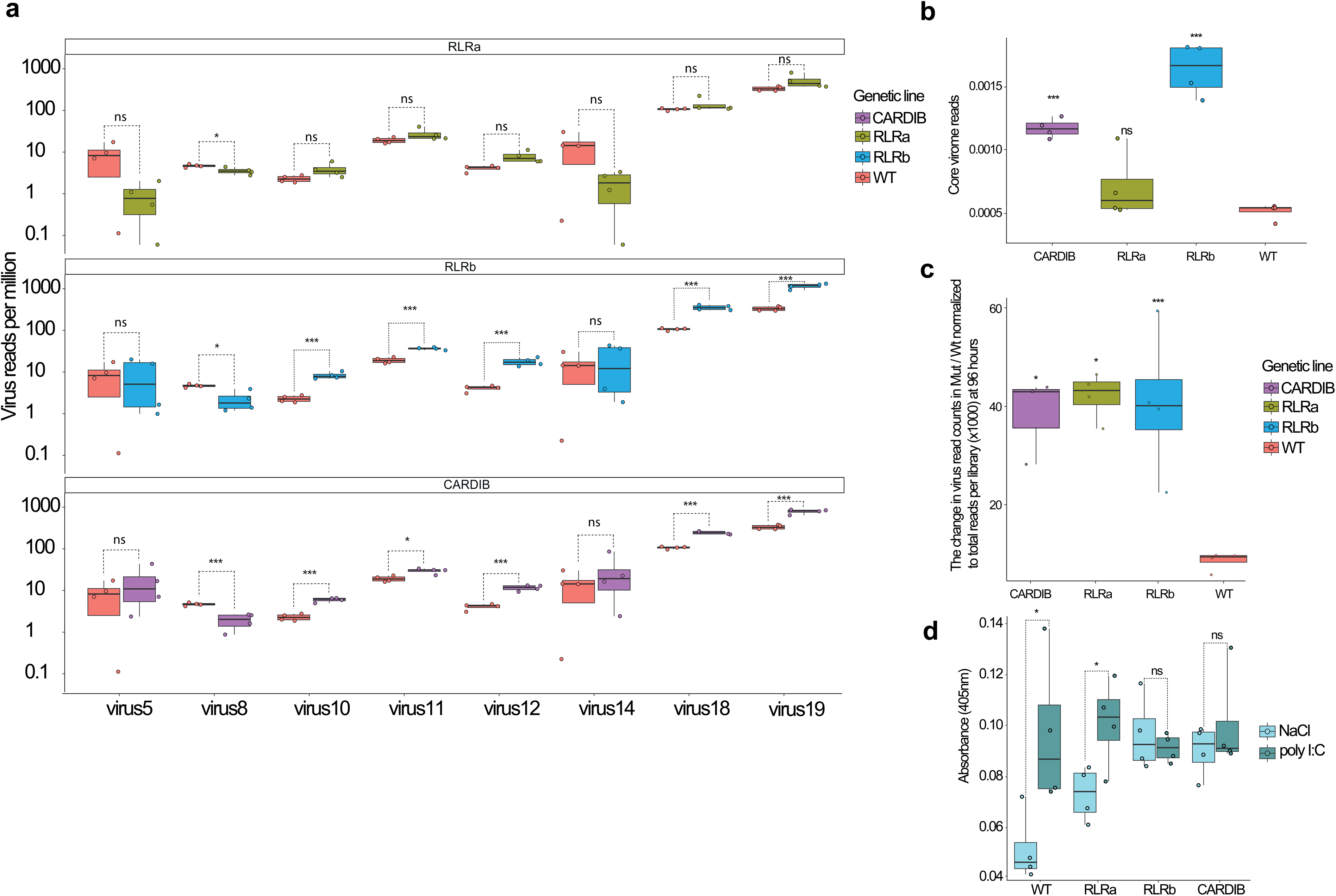
Viral load sequencing-based measurements in wild-type and mutant *N. vectensis* lines. **a,** Change in the viral load of the most abundant viruses in *N. vectensis* core virome under basal lab conditions in adult mutant animals compared to wild-type animals. **b,** Change in the total load of the *N. vectensis* core virome under basal lab conditions in adult mutant animals compared to wild-type animals that were used as a control. The virus read counts were normalized to the total read counts of each sample. Four biological replicates were made for wild-type, *RLRa* KO, and *RLRb* KO. For *CARDIB,* three biological replicates were made **(Supplementary file 1 Tables 19-20)**. **c,** Change of the viral load in mutant and wild-type animals in mesocosm conditions. The animals were exposed to native estuary water inhabited by *N. vectensis* at Georgetown, South Carolina. Viral load was measured at the beginning of the experiment and after 96 h of exposure to estuarine water by sequencing and examining viral read counts **(Supplementary file 1 Tables 11-12 and Supplementary Fig. 5-6)**. The change was measured by comparing the increase in viral load in the mutant lines to that in the wild-type animals after 96 h of exposure to the native water. The virus read counts were normalized to the total read counts of each sample. **d,** the change in the apoptotic activity in the mutant lines compared to wild-type zygotes after injection of poly(I:C). NaCl injection was used as a control. Four biological replicates were made for each condition. Apoptotic activity was measured 24 h post-injection **(Supplementary file 1 Tables 8-10)**. Significance level was tested by paired two-tailed t-test; *p-value < 0.05, **p-value < 0.01, ***p-value < 0.001; NS, not significant.

To evaluate viral susceptibility under natural conditions, sea anemones were transferred from the laboratory to a mesocosm setup and exposed for 96 h to native estuary water directly pumped from a lagoon natively inhabited by *N. vectensis* at Belle W. Baruch Marine Field Laboratory (Georgetown, South Carolina, USA). Viral load was assessed before and after this treatment by sequencing the viromes and examining viral read counts **(Fig. 5c)**. Following mesocosm exposure, all mutant lines showed a significant increase in viral load relative to wild-type after 96 h, confirming the critical role of CARDIB, RLRa and RLRb in host defense under natural conditions.

Finally, we examined the apoptotic response to immune activation, as this is a major line of defense against viruses in *N. vectensis* like in many other animals^23^. Zygotes from mutant and wild-type animals were injected with poly(I:C) or NaCl as a control, and apoptotic activity was measured 24 h post-injection using a Caspase-3 assay kit **(Fig. 5d)**. Wild-type animals showed a strong apoptotic response to poly(I:C) compared to NaCl, while *RLRa* mutants displayed a weaker but still significant differential response. In contrast, *CARDIB* and *RLRb* mutants showed no significant difference in their apoptotic response to poly(I:C) and NaCl. Notably, the apoptotic activity in the *CARDIB* and *RLRb* mutants after NaCl injection is already comparable in its magnitude to that of the wild-type animals injected with poly(I:C) **(Fig. 5d)**.

Overall, these results demonstrate that loss of *CARDIB* or *RLRb* severely compromises the immune system of *N. vectensis*, leading to increased viral load and impaired apoptosis. In contrast, removal of *RLRa* reduces the efficiency of the immune response but does not abolish it. Yet, *RLRa* still seems to be crucial for antiviral immunity under field conditions.

## Discussion

The discovery that *CARDIB* is positioned immediately adjacent to *RLRb* in the *N. vectensis* genome, and that this genomic arrangement is conserved across multiple anthozoan species for at least 648 million years^34^, highlights a striking case of evolutionary stability **(Fig. 1a-b)**. Such long-term conservation strongly suggests that this topology is advantageous, likely reflecting functional interdependence or regulatory coordination between the two genes^35^. Indeed, *CARDIB* and *RLRb* appear to be tightly co-regulated under basal conditions, maintaining immune homeostasis through coordinated expression. However, this co-regulation appears to be disrupted under immune challenge, as shown by their divergent expression patterns following poly(I:C) exposure, a pattern that may reflect a regulatory switch from repression to activation once viral RNA is detected^22^.

This model aligns with the functional data, demonstrating that CARDIB acts as a repressor of immune signaling. The upregulation of immune-related genes in *CARDIB* knockout animals, contrasted with their downregulation upon *CARDIB* overexpression, supports its role as a negative regulator that restrains immune activation under normal conditions **(Fig. 1c-g)**. Such a mechanism may prevent deleterious immune responses in the absence of infection, a key feature of immune balance conserved across the anthozoan evolution.

Structurally, CARDIB retains the canonical CARD domain found in vertebrate MAVS, yet the protein’s overall architecture and function have diverged substantially. Unlike MAVS, which integrates RLR signals to activate downstream interferon responses^36^, CARDIB appears to have undergone “functional inversion”, acting instead to suppress immune activity. This shift suggests that while the CARD domain was evolutionarily conserved as a structural signaling interface, its functional output diversified between cnidarians and vertebrates^37^.

In vertebrates, the RLR proteins RIG-I and MDA5 both contain tandem CARD1-CARD2 domains that activate MAVS through CARD-CARD interactions, yet they differ substantially in their regulation and signaling dynamics. In RIG-I, activation of the 2CARD module depends on K63-linked ubiquitination by TRIM25 and (E3 ubiquitin ligase) Riplet, where CARD2 binds ubiquitin and stabilizes the active tetramer, and CARD1 engages MAVS directly^38–41^. In contrast, MDA5 exposes and oligomerizes its CARDs along long viral dsRNA filaments with minimal ubiquitin dependence, leading to a slower but sustained antiviral response^42,43^. This asymmetry between CARD1 and CARD2, together with divergent activation mechanisms between RIG-I and MDA5, reflects adaptive specialization of CARD signaling interfaces across vertebrates^10^.

Comparative structural analysis of RLRs across metazoans reveals a clear evolutionary transition in domain organization. While vertebrate RIG-I and MDA5, possess two N-terminal CARD domains that mediate downstream signaling through MAVS activation, the *N. vectensis* homologs *RLRa* and *RLRb* retain only a single CARD domain. This change in the number of CARD domains in bilaterians might be caused by the evolutionary pressure from viruses, leading them to evolve their immune signaling to improve their antiviral defense^10,44^. The AlphaFold2 structural predictions reinforce this view, demonstrating that *N. vectensis* RLRs structurally align with vertebrate MDA5 CARD1 but lack a detectable CARD2-like fold, implying that CARD1 is related to an ancestral form of the CARD domain that is conserved across evolution **(Fig. 2a-b)**. At the same time, CARD2 is an addition that has evolved only in bilaterians and enhances the versatility of signaling interactions.

The experimental data support a model in which CARDIB and RLRb interact directly under basal, unstimulated conditions, forming a repressive complex that maintains immune quiescence. Upon introduction of dsRNA to the cytoplasm, RLRb binds it and signals downstream. The strong correlation observed in single-cell RNA-seq (scRNA-seq) data between *CARDIB* and *RLRb* expression under control conditions, together with its marked disruption upon poly(I:C) challenge, provides transcriptional support for the inhibitory role of CARDIB **(Fig. 2c-f)**.

Two independent experimental approaches: proximity ligation assays (PLA) **(Fig. 2g-k),** and Co-IP **(Fig. 2l-m)**, converge to support the physical association between *CARDIB* and *RLRb*. In light of the fact that CARDIB is essentially a CARD domain with barely any additional amino acids, the interaction is most likely direct and CARD-mediated. Hence, CARDIB in anthozoans and MAVS in bilaterians use similar structural means to bind RLRs. However, the regulatory output of this interaction has been inverted: instead of activating downstream signaling, as in the MAVS-RLR system of vertebrates^43,45^, the CARDIB-RLRb interaction serves to restrain it. This supports the notion that cnidarian antiviral signaling represents not a simple immune system, but an alternative evolutionary trajectory built upon ancient CARD-based molecular logic, which is utilized differently.

This evolutionary separation does not preclude molecular resemblance. The observed sequence and structural similarity between *CARDIB* and vertebrate *MAVS* likely represent a slower evolutionary pace, shaped by similar purifying selective pressures to maintain CARD-CARD interaction with RLRs **(Fig. 3a-b)**.

Importantly, these findings highlight the limitations of inferring orthology solely from sequence similarity. Although *CARDIB* shares detectable similarity with *MAVS*, the phylogenetic and experimental evidence together demonstrates that structural resemblance does not equate to shared ancestry or function. Instead, *CARDIB* may represent a parallel evolutionary branch in the evolution of innate antiviral immunity, a molecular prototype that may have evolved alongside, rather than ancestral to, the vertebrate mitochondrial signaling axis^46,47^. Here, we demonstrate how conserved structural domains and rate-dependent attraction can produce the illusion of deep homology and superficially imply common function, an interpretive pitfall increasingly relevant as structural genomics expands cross-kingdom comparisons of immune systems^48,49^.

The functional assays, where *CARDIB*, *RLRa*, and *RLRb* knockout lines were exposed to poly (I:C) provide compelling evidence that both *CARDIB* and *RLRb* are indispensable for initiating the antiviral immune response in *N. vectensis*. Deletion of *RLRb* almost completely abolished the transcriptional response to poly(I:C), underscoring its role as a primary sensor. This finding is consistent with previous evidence showing that RLRb binds long dsRNA, the molecular hallmark of viral infection, whereas RLRa does not^22^. The loss of immune activation in *RLRb*-deficient animals therefore reflects a direct breakdown in the initial viral recognition step required for triggering downstream antiviral programs **(Fig. 4a, d, f)**.

Strikingly, the knockout of *CARDIB* produced a similarly severe phenotype, characterized by the complete absence of immune induction following poly(I:C) stimulation **(Fig. 4c, f)**. This observation reveals that, despite acting as a repressor under basal conditions, *CARDIB* is essential for proper immune activation upon challenge. Such dual functionality, restraining immune gene expression in the absence of infection while permitting or even facilitating activation during stress, illustrates a finely tuned regulatory mechanism that balances immune readiness with homeostasis.

In contrast, *RLRa* knockout animals exhibited only a modest reduction in immune response efficiency, suggesting a supportive or modulatory role rather than a primary sensing function **(Fig. 4b, e, f)**. The presence of a largely intact transcriptional program in *RLRa* mutants indicates that *RLRb* and *CARDIB* form the core antiviral axis in *N. vectensis*, while *RLRa* may contribute to signaling amplification or fine-tuning under specific conditions^22^.

Our results reveal that the CARDIB-RLRb axis is not only central to immune gene activation but also indispensable for maintaining viral control in *N. vectensis*. Under laboratory conditions, which provide optimized and pathogen-reduced environments^33^, knockout lines of either *CARDIB* or *RLRb* resulted in a marked increase in viral load, demonstrating that the loss of either component severely compromises the host’s capacity to suppress viral replication. In contrast, *RLRa* mutants displayed viral loads comparable to wild-type animals, consistent with transcriptomic findings showing only a mild reduction in immune activation **(Fig. 5a-b)**. These observations firmly establish *RLRb* as a primary viral RNA sensor and *CARDIB* as its critical regulatory partner, together forming the functional backbone of the *N. vectensis* antiviral defense system.

Yet, exposure to a mesocosm environment, mimicking natural estuarine conditions, complicates the picture. All three mutant lines (*CARDIB*, *RLRa*, and *RLRb*) exhibited significantly higher viral loads relative to wild-type, probably reflecting the greater viral diversity and more fit pathogens encountered in natural habitats **(Fig. 5c)**. This result underscores that immune components appearing partially redundant under laboratory settings may reveal essential contributions when tested under more complex, ecologically realistic conditions, a phenomenon known as “rewilding”^50^. In particular, the increased viral susceptibility of the *RLRa* mutants in the mesocosm suggests that *RLRa* might act as a secondary responder or enhancer of antiviral defense in vivo, a function that remains hidden in simplified laboratory assays. The phenomenon of rewilding has been documented in mice and zebrafish, where immune phenotypes become evident only under natural exposure conditions, as opposed to laboratory conditions, and display a context-dependent effect^50,51^.

When a host is infected by a virus, one of the harshest tools available to the host to defend itself is the activation of apoptosis^52,53^. The apoptosis assay we performed further supports a dual role for *CARDIB* and *RLRb* in regulating both antiviral signaling and cell death pathways. Wild-type animals showed significant, stimulus-dependent apoptotic activation following poly(I:C) injection, whereas *CARDIB* and *RLRb* mutants failed to exhibit such activation. Instead, they displayed elevated basal apoptotic activity, suggesting loss of proper regulation control over immune-linked cell death. Interestingly, *RLRa* mutants exhibited an intermediate phenotype of a weaker yet still significant apoptotic response to poly(I:C) **(Fig. 5d)**. This partial activity indicates that while *RLRa* is not essential for initiating apoptosis, it likely contributes to amplifying or sustaining the response once immune activation has been triggered. Such a modulatory role may serve to fine-tune antiviral signaling and prevent either insufficient or excessive cell death, helping to maintain balanced immune homeostasis under stress conditions. The chronic activation state observed in the *CARDIB* and *RLRb* mutants may reflect persistent immune dysregulation resulting from the loss of CARDIB-mediated repression and RLRb-dependent sensing, alternatively, an increase in viral load that leads via other ligands and receptors to immune response and apoptosis.

Collectively, these findings demonstrate that disruption of either *CARDIB* or *RLRb* leads to a dual failure: the inability to mount an effective antiviral response and a simultaneous breakdown of apoptotic regulation. The resulting elevation in viral burden and loss of homeostasis highlights the integrated nature of antiviral defense and immune regulation in Anthozoa. This interplay between sensing viruses and suppression of immunity mediated through various CARD-based interactions suggests that dynamic balancing of activation and repression is an ancient and fundamental necessity stemming back to the last common ancestor of the Cnidaria and the Bilateria, and possibly much earlier in evolution^54^.

## Materials and Methods

### Sea Anemone Culture

Early life stages of *N. vectensis* (embryos, larvae, and primary polyps) were maintained in the dark at 22 °C in artificial seawater with a salinity of 16 ‰, whereas juveniles were reared at 18 °C. Starting two weeks post-fertilization, polyps were fed *Artemia salina* nauplii three times per week. Gamete spawning was induced as previously described^55^. For microinjection experiments, fertilized eggs were dissociated from the egg package by incubation in 3% L-cysteine (Merck Millipore, USA). For all other purposes, fertilized eggs were kept at 22 °C for three days until planula larvae were spontaneously released from the gelatinous egg mass. All *N. vectensis* individuals used in this study belonged to the common laboratory strain originating from the Rhode River, Maryland, USA^56^.

### Injection of viral mimics to wild-type and mutants

To activate the antiviral immune response in *N. vectensis*, we used dsRNA as a viral RNA mimic. We applied 6.25 ng/ml of high molecular weight (HMW) poly(I:C) (Invivogen, USA) in 0.9 % NaCl, with an average size of 1.5-8 kb, and used 0.9 % NaCl as the control. This poly(I:C) concentration was selected after testing a range of doses, as it proved the most effective for assessing the antiviral response. Higher concentrations resulted in high mortality and abnormal zygote morphology within 24 h. In each experiment, 150-600 zygotes were injected per group and maintained at 22 °C^22^.

### Generation of knockout mutant lines of RLRa, RLRb and CARDIB

CRISPR/Cas9 genome editing in *N. vectensis* embryos was performed following an established protocol^57^. For each gene, two gRNAs were designed using the CRISPOR web tool^58^. Each sgRNA targeting exons was ordered from Integrated DNA Technologies Inc (IDT, USA), using oligos described in **Supplementary file 1 Table 3**. The injection mix was prepared at room temperature and incubated for 10 min before injection into fertilized eggs, contained recombinant Cas9 protein with NLS (1500 ng/μl; PNA Bio, USA; CP0120) and sgRNA (750 ng/μl).

Genomic DNA from F0 primary polyps was extracted by washing samples three times with 100 % MeOH in PCR tubes. Samples were dried at 50 °C for 10 min, after which DNA extraction buffer (10 mM Tris pH 8, 1 mM EDTA pH 8, 25 mM NaCl, and 200 μg/ml Proteinase K; Thermo Fisher Scientific, USA) was added. Samples were incubated for 2 h at 50 °C, followed by 5 min at 96 °C to inactivate Proteinase K. Mutation analysis was performed using high-resolution melting (HRM) on a Magnetic Induction Cycler qPCR instrument (Bio Molecular Systems, Australia) to detect indels. PCR fragments adjacent to the targeted regions were amplified, with primers listed in the **Supplementary file 1 Table 4**. F0 injected animals were reared to sexual maturity (>4 months) and induced to spawn gametes following an established protocol^59^. F0 mutants were crossed with wild-type individuals to generate F1 heterozygotes. Genomic DNA was extracted from the progeny of each tentative F0 founder. Mutation detection was carried out in F0 animals, and F1 carriers were raised to sexual maturity (>4 months). One F1 heterozygous individual was crossed with another F1 heterozygote carrying the same mutation, and F2 progeny from this cross were raised to 2 months of age. These F2 individuals were placed in 12 well plates and individually genotyped to identify wild-type, heterozygous, and homozygous null individuals.

Tentacle tissue biopsies were collected using a scalpel, with tissue washed five times in 100% MeOH prior to DNA extraction (as for F0 animals). After identifying homozygous knockout mutants, they were crossed with other F2 individuals to produce F3 homozygous mutants for downstream experiments^59^.

### mRNA synthesis

For mRNA synthesis, we followed a previously described method^60^. For injection into *N. vectensis* zygotes, the gBlock template (IDT) was amplified for use as an mRNA template. mRNA was transcribed with the HiScribe T7 mRNA Kit with CleanCap Reagent AG (New England Biolabs, USA) according to the manufacturer’s protocol, using 1 µg of amplified template. *In vitro* transcribed products were purified with the RNA Clean & Concentrator 25 kit (Zymo Research, USA) and eluted in 33 µl of RNase-free water. Concentrations were measured with the Qubit RNA Broad Range Assay Kit and Qubit Fluorometer (Thermo Fisher Scientific).

Polyadenylation was performed with *E. coli* Poly(A) Polymerase (New England Biolabs) for 30 min at 37 °C, after which products were further purified with the RNA Clean & Concentrator 5 kit (Zymo Research) and eluted in 8-10 µl of RNase-free water. The integrity of the single product was validated on a 1.5% agarose gel after incubation at 95 °C for 2 min in a thermocycler with a heated lid, followed by cooling to 22 °C and mixing with formamide (Merck Millipore) in a 1:3 ratio. The mRNA was stored at -80 °C until injection. mRNA templates and primers used for cloning and amplification are listed in **Supplementary Fig. 2**.

### Cell dissociation

Embryos were dissociated into single cells as previously described^61^. Briefly, individuals were washed twice with calcium/magnesium free and EDTA free artificial seawater (×3 stock solution is: 17 mM Tris-HCl, 165 mM NaCl, 3.3 mM KCl and 9 mM of NaHCO_3_; final solution pH 8.0) and incubated with 50 µg/ml liberaseTM (Roche, Switzerland) at room temperature for 5-10 min with occasional gentle pipetting, until fully dissociated. The reaction was stopped by adding 1/10 volume of 0.5 M EDTA solution. The suspension was filtered using a 35 µm cell strainer (Jet Bio Filtration, China). Cells were then centrifuged at 500 × g at 4 °C and resuspended in 1× calcium/magnesium-free sterile PBS (Hylabs, Israel) supplemented with 2 % BSA (MP Biomedicals, USA). Cells were counted on a hemocytometer and viability was determined using DAPI (Thermo Fisher Scientific).

### Proximity ligation assay

Cells derived from 24 h old planulae after injection with mRNA encoding either CARDIB-FLAG-GFP or GFP alone (control) were dissociated as described above. Dissociated cells were then washed twice with PBS. Cells were fixed with 4 % paraformaldehyde (PFA) in PTW (PBS, 0.1 % Tween-20) for 30 min on a rotator at 4 °C. In the following steps, centrifugation was performed for 3 min at 3000 x g at 4 °C. Cells were resuspended in wash buffer (PBS, 0.2 % BSA, 0.1 % Triton-x), washed twice, and counted using a 1:10 concentration of DAPI (Thermo Fisher Scientific) on a hemocytometer. The samples were blocked for 3 h in a solution containing 20 % sheep serum (Merck Millipore) and 3 % bovine serum albumin (BSA, fraction V; MP Biomedicals) in PBS with 0.1 % Triton X-100. Primary antibodies against RLRb and against FLAG (GenScript, USA) were used at 1:1000 and 1 µg/ml, respectively. This concentration was determined based on titration experiments that demonstrated a decent signal-to-noise ratio. After incubating overnight at 4°C, cells were then washed twice in Wash Buffer (PBS, 0.2 % BSA, 0.1 % Triton-X) and then in Wash Buffer A (Merck Millipore). Following the steps of the proximity ligation assay was performed according to the manual instructions of PLA Duolink kit (Merck Millipore). The samples were incubated with 40 μl reaction mixture (8 μl PLA probe MINUS stock, 8 μl PLA probe PLUS stock, and 24 μl PBS) in a chamber for 1 h at 37 °C. The samples were washed in 1x Wash Buffer A for 2×10 min on the shaker at RT, and ligation was performed in a 40 μl reaction: 1 μl of ligase to 39 μl of ligation solution. Samples were incubated in the ligation reaction mixture for 30 min at 37 °C, then washed 2×10 min in Wash Buffer A on the shaker at RT. 40 μl of amplification reaction (0.5 μl polymerase and 39.5 μl amplification solution) was added to each sample before incubation at 37 °C for 100 min. Next, the cells were stained with 1:250 DAPI for nuclei staining and were washed in Wash Buffer B (Merck Millipore) for 2×10 min on the shaker at RT, and in 0.01 Wash Buffer B for 2 min. The samples were mounted with Vectashield Mounting Medium (Vector Laboratories, USA). An inverted confocal microscope (A1R by Nikon, Japan) was used for sample visualization. Images were processed using Fiji^62^ and Photoshop (Adobe, USA) for channel merging. Processing of raw photographs was performed evenly on all files, and no manipulations were made to enhance the uneven part of any picture.

### Western blot

Wild-type, knockout mutants, CARDIB-overexpressing, and poly(I:C)/NaCl-injected animals were flash frozen in liquid nitrogen and homogenized with an electric pestle in lysis buffer (50 mM Tris-HCl pH 7.4, 150 mM KCl, 0. 5% NP-40, 10 % glycerol, and inhibitor Cocktail Set III, EDTA-free by Merck Millipore). Lysates were incubated at 4 °C for 2 h with rotation and then centrifuged at 16,000 × g for 10 min at 4 °C. The supernatant was collected, and protein concentration was measured using the BCA Protein Assay Kit (Cyanagen, Italy). Samples were normalized to equal protein concentrations, mixed with blue loading buffer (New England Biolabs) at a 1:2 ratio, and boiled at 100 °C for 8 min.

Equal amounts of protein were separated on 4-15 % Mini-PROTEAN TGX Precast Protein Gels (Bio-Rad, USA) and transferred to nitrocellulose membranes (Bio-Rad). Membranes were washed with TBST buffer (20 mM Tris pH 7.6, 150 mM NaCl, 0.1% Tween-20) and blocked with 5 % skim milk (BD, USA) in TBST for 1 h at room temperature on a shaker. Membranes were then incubated overnight at 4 °C with either target polyclonal antibodies or monoclonal mouse anti-GAPDH antibody (GA1R, ab125247; Abcam, UK) diluted 1:1000 in TBST containing 5 % BSA (MP Biomedicals, USA) inside sealed sterile plastic bags.

After three 10 min washes in TBST, membranes were incubated for 1 h with peroxidase-conjugated secondary antibodies (anti-rabbit, anti-mouse, or anti-Guinea pig; Jackson ImmunoResearch, USA) diluted 1:10,000 in 5 % skim milk in TBST. Membranes were washed three times with TBST, and detection was carried out using the Clarity ECL or Clarity Max kits (Bio-Rad) according to the manufacturer’s instructions. Chemiluminescent signals were visualized with a CCD camera of the Odyssey Fc imaging system (Li-COR Biosciences, USA). Protein size determination was performed by simultaneously running the Precision Plus Protein Dual Color Protein Ladder (Bio-Rad) and scanning at 700 nm. Band intensities were quantified with Image Studio software (Li-COR Biosciences), and fold changes were calculated relative to the loading control for each sample. All original western figures and band intensity measurements are available in the link:10.6084/m9.figshare.30105067, and **Supplementary Fig. 3**.

### CARDIB and RLRb co-immunoprecipitation

Protein A Magnetic Beads (MedChemExpress, USA) were washed five times in 1 ml of 1× PBS. Antibodies against RLRb (5 µg; GeneScript)^22^ or anti-NeNaC2 (5 µg; GeneScript)^63^ were added to the beads in 1.4 ml of PBS. Samples were rotated overnight at 4 °C.

24 h old *N. vectensis* embryos were dissociated into single cells with Liberase TM (Roche, Switzerland) as previously described^23,61^. Cells were then cross-linked using dithiobis succinimidyl propionate (DSP; Cayman Chemicals, USA) as previously described^64^. Briefly, dissociated cells were washed twice with PBS and incubated with DSP at a final concentration of 0.8 mg/ml for 30 min at 4 °C with rotation. Cross-linking was quenched by adding 0.2 M glycine for 15 min at 4 °C. Cells were washed with PBS and lysed for 30 min at 4 °C in lysis buffer (50 mM Tris-HCl, pH 8, 150 mM NaCl, 1% NP-40, protease inhibitor cOmplete ULTRA tablets [Roche], and protease inhibitor Cocktail Set III, EDTA-free [Merck Millipore]). Samples were centrifuged at 16,000 × g for 10 min at 4 °C, and the supernatant was collected. Protein concentration was determined using the BCA Protein Assay Kit (Cyanagen), and lysates were normalized to equal protein amounts.

For pre-clearing, 100 µl of Protein A Magnetic Beads (MedChemExpress) were washed three times with 1 ml of PBS, and the tissue lysate was added to the beads. Lysis buffer was added to a final volume of 1.4 ml, and samples were rotated for 1 h at 4 °C. After discarding the beads, pre-cleared lysates were then incubated with the antibody-bound beads for 2 h at 4 °C with rotation. After incubation, lysates were discarded, and the beads were washed twice with PBS. Beads were then resuspended in 60 µl of double-distilled water and 30 µl of blue loading buffer (New England Biolabs).

Co-IP samples were boiled at 100 °C for 8 min and centrifuged at maximum speed at 4 °C. The beads were discarded, and the supernatants were collected for western blot analysis as described above.

### Analysis of RLRs and CARDIB genomic structure

To reconstruct the evolution and conservation of microsynteny between CARDIB and RLRs, we employed a comparative genomic approach across anthozoan genomes. The genomes analyzed included *Xenia sp.*^65^, *Acropora muricata*^66^, *Exaiptasia diaphana*^67^, and *N. vectensis*^68^.

Using *N. vectensis* CARDIB and RLRb protein sequences as queries, we performed tBLASTn searches (e-value threshold: 1e-5) to identify genomic locations and gene models of *CARDIB* and *RLR* orthologs in the other anthozoan genomes. In *Xenia*, gene models were fragmented; therefore, we conducted an additional de novo transcriptome assembly to better resolve gene structures. Raw sequencing reads were downloaded from the NCBI Sequence Read Archive (PRJNA548325) and assembled using Trinity v2.15.2. BLASTx searches were then carried out using the assembled contigs as queries against *N. vectensis* CARDIB and RLRb protein sequences. Subsequently, BLASTn searches were performed on the identified CARDIB and RLRb contigs to refine gene models.

To classify anthozoan sequences as either RLRa or RLRb, the protein sequences identified in this study were added to a previously published RLR alignment used for phylogenetic analyses^22^. The combined dataset was used to reconstruct the evolutionary history of RLRs. Protein sequences were aligned using MUSCLE implemented in MEGA 11^69^. The alignments were then imported into IQ-TREE, and the best-fit model of protein evolution was determined using ModelFinder^70^. Based on the Bayesian information criterion, the WAG+I+G4 model was selected. Phylogenetic trees were inferred from the alignments using 1,000 ultrafast bootstrap replicates and the SH-aLRT test^71^. Sequences clustering with *N. vectensis* RLRs were subsequently used to define sequence orthology **(Supplementary Fig. 4)**.

### RNA-seq library preparation and sequencing

RNA of the samples from the injected animals was extracted with Quick-RNA Microprep kit (Zymo Research, USA) according to the manufacturer’s protocol, treated with 4 ml of Turbo DNase (Thermo Fisher Scientific), and re-extracted with the Quick-RNA Microprep kit (Zymo Research). The quality of total RNA was assessed on Bioanalyzer Nanochip (Agilent, USA), and only samples with RNA integrity number >7.0 were retained. For the generation of the ployA-selected RNA-seq libraries, we used the service of Novogene (Singapore) in their facilities according to their in-house protocol and sequenced on NovaSeq 6000 (Illumina, USA) using paired-end 150 bp chemistry.

RNA of the samples from the adult animals was extracted by flash freezing of the samples, grinding them with mortar and pestle and then the resulting powder was extracted with Trizol (Thermo Fisher Scientific) according to the manufacturer’s protocol, treated with 2 ml of Turbo DNase (Thermo Fisher Scientific), and re-extracted again with Trizol. The quality of total RNA was assessed on Bioanalyzer Nanochip (Agilent), and only samples with RNA integrity number >7.0 were retained.

RNA-seq libraries from adults were generated using Zymo-Seq RiboFree Total RNA Library Kit (Zymo Research) following the manufacturer’s protocol and sequenced on NextSeq 2000 (Illumina) using single-end 100 bp chemistry at the Genomic Technologies Center of the Hebrew University of Jerusalem.

### RNA-seq analysis

The quality of raw sequencing reads was assessed and visualized using FastQC (S. Andrews. Babraham Bioinformatics, Babraham Institute, 2010.). Adapter trimming and quality filtering were performed with Trimmomatic (v0.36)^72^ using the parameters: ILLUMINACLIP:TruSeq3-SE:2:30:10, LEADING:3, TRAILING:3, SLIDINGWINDOW:4:15, and MINLEN:36. Cleaned reads were aligned to the *N. vectensis* reference genome^73^ using STAR (version 2.7.10a)^74^ with default alignment parameters. Gene-level counts were generated with featureCounts (v2.0.1)^75^. Differential expression analysis was conducted using DESeq2^76^ and EdgeR^77^. Genes with an absolute log_2_ fold change > 1 and an adjusted p-value < 0.05 were defined as differentially expressed genes (DEGs). Variance-stabilizing transformation (VST) of read counts was applied using the DESeq2 vst function (blind = FALSE), and principal component analysis (PCA) was performed on the transformed data. VennDiagram^78^ was used to generate Venn diagrams. Over-representation analysis (ORA) of DEGs was performed using clusterProfiler^79^. Gene Ontology (GO) annotations were retrieved from the QuickGO database^80^.

### Clustering and phylogenetic analysis of CARD domains

A total of 1,055 CARD domain sequences were retrieved from 859 UniProt entries, representing six major metazoan groups (Cephalochordata, Cnidaria, Echinodermata, Protostomia, Urochordata and Vertebrata) **(Supplementary file 1 Table 5)**. A clustering analysis of the sequences was carried out using CLANS software^81^ with default settings and 300,000 rounds. The 2D coordinates generated by CLANS were subsequently analyzed in R to determine the optimal number of clusters. We applied the Elbow, Silhouette, and Gap statistic methods via the factoextra package^82^ to evaluate clustering performance, and k-means clustering was then performed with five clusters (k = 5) using 25 random initializations. A phylogenetic analysis was conducted on the same set of sequences. The 1,055 sequences were aligned using MAFFT v7^83^, and the resulting alignment served as input for IQ-TREE2^84^ to infer a maximum-likelihood tree. Node support was assessed using three statistical measures: SH-aLRT, aBayes, and bootstrap.

### Evolutionary and selection analyses of CARD domains

We selected 90 representative CARD-domain protein sequences, 15 from each of the six well-supported clades identified in the previously reconstructed CARD phylogeny **(Fig. 3a)** and extracted the corresponding subtree for subsequent analyses. Amino acid alignments were generated using MAFFT v7.520 9^85^ with the L-INS-i algorithm, codon alignments were produced with PAL2NAL v14 to preserve reading frames and codon positions. The final dataset comprised 90 sequences and 106 codons. We performed likelihood-based codon model analyses in PAML v4.10.7 (codeml)^86^ to assess variation in selective pressure as defined by ω (i.e., dN/dS) among lineages and sites. For the branch model (two-ratio test), we compared a null model assuming a single ω across all branches with an alternative model assigning distinct ω values to the foreground branches (MAVS clade and the *N. vectensis* CARDIB clade) and the remaining background branches (the four other well-supported clades). For the branch-site model (model A), we tested positive selection acting on a subset of sites along the foreground clades. The null model fixed ω_₂_ = 1, whereas the alternative model estimated ω_₂_ freely. Model comparisons were conducted using likelihood ratio tests (LRTs) evaluated against a χ² distribution with 1 degree of freedom. Additional details regarding the foreground annotation of the branches can be found in **Supplementary file 2.**

### In silico protein structure prediction

The protein structures of *NveRLRa* and *NveRLRb* were predicted using AlphaFold2^87^. The MDA5 CARD domains (PDB: 7DNI) were superimposed onto the N-terminal domains (NTDs) of *NveRLRa* and *NveRLRb*, which include a CARD domain and an adjacent uncharacterized region preceding the helicase domain, using PyMOL (The PyMOL Molecular Graphics System, Version 3.0 Schrödinger, LLC). Structural similarity was assessed by calculating the Root Mean Square Deviation (RMSD).

### Gene co-expression analysis across conditions using single-cell transcriptomic data

We analyzed single-cell transcriptomic data generated in our laboratory^30^ to examine the relationship between the expression levels of *RLRb* and *CARDIB*, measured as UMI fractions per metacell, across three experimental conditions (non-injected, NaCl-injected, and poly(I:C)-injected animals) **(Supplementary file 1 Table 6)**. Pearson correlation coefficients were computed in R using cor.test (method = ’pearson’), both (i) across all metacell-condition averages combined and (ii) separately within each experimental condition. For each correlation, we report Pearson’s r along with the corresponding p-value.

### Antibody generation

Custom polyclonal antibodies were raised against recombinant fragment antigens generated by rabbit host immunization (GenScript, USA and ProteoGenix, France). Each recombinant fragment was injected into three rabbit hosts. Sera after 3rd immunization was used for ELISA and western blot verification, followed by affinity purification against the recombinant antigen **Supplementary Fig. 5**). Amino acid sequences of antigen fragments used for antibodies generation are presented in **Supplementary file 1 Table 7**.

### Apoptotic reactivity assay

To identify apoptotic reactivity, we followed the method as previously described^23^. For each condition (eGFP vs. FLAG-tagged CARDIB), about 150 injected animals were snap frozen in liquid nitrogen and stored in -80°C. Caspase-3 Assay Kit (colorimetric) (Abbkine, USA) was used according to the manufacturer’s instructions. The embryo pellet was lysed in cell lysis buffer using a homogenizer. Protein yield was measured by an Epoch Microplate Spectrophotometer (BioTek Instruments, USA) equipped with a Take3 Plate. After measurement, equal protein amounts were loaded into each reaction (21.385mg). Absorbance was measured at 405 nm after 2 h of incubation at 37 °C. For detailed parameters, tables are shown in **Supplementary file 1 Tables 8-10**.

### Mesocosm field survival assay

Prior to the experiment, all animals were starved for two days and acclimated from the standard laboratory salinity of 15 ppt to 30 ppt. This adjustment was made to reduce the likelihood of detecting effects related to salinity stress, as natural salinity at the study site is typically close to full-strength seawater.

Mesocosms were deployed in August 2023 at the Belle W. Baruch Marine Field Laboratory (Georgetown, South Carolina, USA) **(Supplementary Fig. 6a-b)**. A total of five replicate mesocosms were constructed. Each mesocosm consisted of a plastic bin (58.5 cm L × 38.1 cm W × 33 cm H) containing a raised tray (39.4 cm L × 25.4 cm W × 15.2 cm H), on which animal containers were placed **(Supplementary Fig. 6b, d, e)**. Each bin was filled with natural estuary water collected from an adjacent salt panne, which was bag-filtered through a 200 µm filter (Pentair Industrial, KO200K2S) to remove debris. Water circulation in each mesocosm was maintained with a submersible pump (30 W, 550 GPH; Freesea, FS-030R) to create a gentle flow.

Animal containers (8.9 cm L × 8.9 cm W × 6.35 cm H) were modified by removing a 5 cm hole from each wall to allow water flow **(Supplementary Fig. 6c-e)**. Animal containers were weighed down with coarsely filtered sediment (Onyx Sand; Seachem, item 3505) and lined internally with 100 µm mesh lining (uxcell, item a24030200ux0303) secured with wooden clothespins. Each container housed a single *N. vectensis* strain with six individuals. Bag-filtered estuary water was added so that the waterline submerged half the container height **(Supplementary Fig. 6c, e)**, which prevented the escape of animals.

Mesocosm water was replaced every two days (48 h) with fresh bag-filtered estuary water. Animals were collected at two timepoints: the initial timepoint before the start of the experiment (T0) and after 96 h (T96). Collected animals were flash frozen and stored at -80 °C. Samples were transported to the UNC Charlotte laboratory on dry ice and immediately stored at -80 °C until extraction.

### N. vectensis virome sequencing and read processing

RNA was extracted from whole animals using the Qiagen All-Prep kit (Qiagen, USA; cat. no. 80204) with minor modifications to the manufacturer’s protocol. In step 2, 600 µl of RLT Plus buffer was added, and animals were homogenized with a 3 ml syringe (BD Plastipak, 309577). Samples were placed on ice for 10 min and then centrifuged as described in step 4 of the protocol. For total RNA purification, 350 µl of 70 % ethanol was added, and the remaining steps were performed according to the manufacturer’s instructions. RNA quantity and quality were assessed with the Qubit High Sensitivity RNA kit (Thermo Fisher Scientific; Q32855), NanoDrop, and TapeStation High Sensitivity RNA kit (Agilent; 5067-5579). Libraries were prepared from 500 ng of total RNA using the Tecan Universal Plus Total RNA-Seq Library Kit with an *N. vectensis* rRNA depletion step (Ref. 0361-24, 0370-24, S02713) and sequenced on a NovaSeq X Plus 10B (Admera Health, USA).

To identify sequence reads originating from viruses and/or microbial taxa, we developed a custom bioinformatics pipeline **(Supplementary Fig. 7)**. Raw reads were first processed with Trimmomatic^72^ and FastQC^88^ to remove adapters and assess quality. Trimmed reads were then aligned to the *N. vectensis* genome^89^ using HISAT2^90^ to filter out host reads. Unmapped reads were extracted with SAMtools^91^ to generate fastq files. Additional rounds of mapping to the *N. vectensis* genome were performed with HISAT2, followed by Bowtie2 using the (--very-sensitive-local parameter)^92^, until no reads mapped to the host genome.

Unmapped reads were further filtered against an rRNA database (SILVA LSU and SSU, GTDB SSU, and Rfam)^93–95^ to remove eukaryotic and prokaryotic rRNA sequences. The resulting dataset, depleted of host and rRNA reads, is hereafter referred to as “presumably viral reads.” To normalize viral read counts, two assemblies were generated with SPAdes rnaviral^96^: the Total-T0 assembly, using all presumably viral reads from 16 T0 samples, and the Total-T96 assembly, using all presumably viral reads from 15 T96 samples (four replicates for each strain at each timepoint, except for CARDIB T96, which had three replicates). Reads from each sample were mapped back to the corresponding assembly using HISAT2, and normalization factors were calculated by dividing the number of mapped viral reads to the Total-T0 or Total-T96 assembly by the number of raw sequencing reads for that sample and multiplying by 1000. Statistical analyses are described in the **Supplementary file 1 tables 11-12**.

## Supporting information

Supplementary Alignments

## Data availability

All sequencing data generated in this work are publicly available under accessions PRJNA1250240 and PRJNA1262874 at the SRA database of the National Center for Biotechnology Information (NCBI) under BioProject: An ancient anthozoan protein reveals an alternative evolutionary path of antiviral signaling and BioProject: *Nematostella vectensis* Total RNA-seq from a mesocosm study conducted in a South Carolina estuary. Bioinformatic processing scripts can be found on GitHub: https://github.com/sydneybirch/Nematostella_virome_Mesocosm_2023, and https://github.com/adrianjaimes/Cnidarian-immune-system

## Acknowledgements

The authors would like to thank Dr. Michal Bronstein and Mrs. Adi Turjeman of the Genomic Technologies Center of the Alexander Silberman Institute of Life Sciences, The Hebrew University of Jerusalem, for their help with sequencing. This work was supported by a European Research Council Consolidator Grant (AntiViralEvo; 863809) and a US-Israel Binational Science Foundation joint Grant with the National Science Foundation to YM and AMR (2020669).

## References

1. Koonin, E. V. & Dolja, V. V. A virocentric perspective on the evolution of life. Curr. Opin. Virol. 3, 546–557 (2013).

2. Tenthorey, J. L., Emerman, M. & Malik, H. S. Evolutionary Landscapes of Host-Virus Arms Races. Annu. Rev. Immunol. 40, 271–294 (2022).

3. Broecker, F. & Moelling, K. What viruses tell us about evolution and immunity: beyond Darwin? Ann. N. Y. Acad. Sci. 1447, 53–68 (2019).

4. Yoneyama, M. et al. Shared and Unique Functions of the DExD/H-Box Helicases RIG-I, MDA5, and LGP2 in Antiviral Innate Immunity. J. Immunol. 175, 2851–2858 (2005).

5. Kang, D. et al. mda-5: An interferon-inducible putative RNA helicase with double-stranded RNA-dependent ATPase activity and melanoma growth-suppressive properties. Proc. Natl. Acad. Sci. U. S. A. 99, 637–642 (2002).

6. Luo, D. et al. Structural insights into RNA recognition by RIG-I. Cell 147, 409–422 (2011).

7. Wu, B. & Hur, S. How RIG-I like receptors activate MAVS. Curr. Opin. Virol. 12, 91–98 (2015).

8. Hou, F. et al. MAVS forms functional prion-like aggregates to activate and propagate antiviral innate immune response. Cell 146, 448–461 (2011).

9. Zeng, W. et al. Reconstitution of the RIG-I pathway reveals a signaling role of unanchored polyubiquitin chains in innate immunity. Cell 141, 315–330 (2010).

10. Mukherjee, K., Korithoski, B. & Kolaczkowski, B. Ancient Origins of Vertebrate-Specific Innate Antiviral Immunity. Mol. Biol. Evol. 31, 140–153 (2014).

11. Seth, R. B., Sun, L., Ea, C.-K. & Chen, Z. J. Identification and Characterization of MAVS, a Mitochondrial Antiviral Signaling Protein that Activates NF-κB and IRF3. Cell 122, 669–682 (2005).

12. Korithoski, B. et al. Evolution of a Novel Antiviral Immune-Signaling Interaction by Partial-Gene Duplication. PLOS ONE 10, e0137276 (2015).

13. Li, X.-D., Sun, L., Seth, R. B., Pineda, G. & Chen, Z. J. Hepatitis C virus protease NS3/4A cleaves mitochondrial antiviral signaling protein off the mitochondria to evade innate immunity. Proc. Natl. Acad. Sci. 102, 17717–17722 (2005).

14. Wang, B. et al. Enterovirus 71 Protease 2Apro Targets MAVS to Inhibit Anti-Viral Type I Interferon Responses. PLOS Pathog. 9, e1003231 (2013).

15. Ding, S. et al. Rotavirus VP3 targets MAVS for degradation to inhibit type III interferon expression in intestinal epithelial cells. eLife 7, e39494 (2018).

16. Sharma, A., Kontodimas, K. & Bosmann, M. The MAVS Immune Recognition Pathway in Viral Infection and Sepsis. Antioxid. Redox Signal. 35, 1376–1392 (2021).

17. Layden, M. J., Rentzsch, F. & Röttinger, E. The rise of the starlet sea anemone Nematostella vectensis as a model system to investigate development and regeneration. Wiley Interdiscip. Rev. Dev. Biol. 5, 408–428 (2016).

18. Technau, U. & Steele, R. E. Evolutionary crossroads in developmental biology: Cnidaria. Development 138, 1447–1458 (2011).

19. 19. Al-Shaer, L., Havrilak, J. & Layden, M. J. Nematostella vectensis as a Model System. in Handbook of Marine Model Organisms in Experimental Biology (CRC Press, 2021).

20. Ikmi, A., McKinney, S. A., Delventhal, K. M. & Gibson, M. C. TALEN and CRISPR/Cas9-mediated genome editing in the early-branching metazoan Nematostella vectensis. Nat. Commun. 5, 5486 (2014).

21. Fraune, S., Forêt, S. & Reitzel, A. M. Using Nematostella vectensis to Study the Interactions between Genome, Epigenome, and Bacteria in a Changing Environment. Front. Mar. Sci. 3, (2016).

22. Lewandowska, M., Sharoni, T., Admoni, Y., Aharoni, R. & Moran, Y. Functional Characterization of the Cnidarian Antiviral Immune Response Reveals Ancestral Complexity. Mol. Biol. Evol. 38, 4546–4561 (2021).

23. Kozlovski, I. et al. Induction of apoptosis by double-stranded RNA was present in the last common ancestor of cnidarian and bilaterian animals. PLOS Pathog. 20, e1012320 (2024).

24. Margolis, S. R. et al. The cyclic dinucleotide 2’3’-cGAMP induces a broad antibacterial and antiviral response in the sea anemone Nematostella vectensis. Proc. Natl. Acad. Sci. U. S. A. 118, e2109022118 (2021).

25. Goubau, D., Deddouche, S. & Reis e Sousa, C. Cytosolic sensing of viruses. Immunity 38, 855–869 (2013).

26. Shaner, N. C. et al. Improved monomeric red, orange and yellow fluorescent proteins derived from Discosoma sp. red fluorescent protein. Nat. Biotechnol. 22, 1567–1572 (2004).

27. Kim, J. H. et al. High Cleavage Efficiency of a 2A Peptide Derived from Porcine Teschovirus-1 in Human Cell Lines, Zebrafish and Mice. PLOS ONE 6, e18556 (2011).

28. Li, K. et al. Insights into the structure and RNA-binding specificity of Caenorhabditis elegans Dicer-related helicase 3 (DRH-3). Nucleic Acids Res. 49, 9978–9991 (2021).

29. Batachari, L. E., Dai, A. Y. & Troemel, E. R. Caenorhabditis elegans RIG-I-like receptor DRH-1 signals via CARDs to activate antiviral immunity in intestinal cells. Proc. Natl. Acad. Sci. 121, e2402126121 (2024).

30. Kozlovski, I. et al. Functional characterization of specialized immune cells in a cnidarian reveals an ancestral antiviral program. 2025.01.24.634691 Preprint at 10.1101/2025.01.24.634691 (2025).

31. Sharoni, T. et al. Heat Stress Drives Rapid Viral and Antiviral Innate Immunity Activation in Hexacorallia. Mol. Ecol. n/a, e70098.

32. Koonin, E. V. & Aravind, L. Origin and evolution of eukaryotic apoptosis: the bacterial connection. Cell Death Differ. 9, 394–404 (2002).

33. Lewandowska, M., Hazan, Y. & Moran, Y. Initial Virome Characterization of the Common Cnidarian Lab Model Nematostella vectensis. Viruses 12, 218 (2020).

34. McFadden, C. S. et al. Phylogenomics, Origin, and Diversification of Anthozoans (Phylum Cnidaria). Syst. Biol. 70, 635–647 (2021).

35. Irimia, M. et al. Extensive conservation of ancient microsynteny across metazoans due to cis-regulatory constraints. Genome Res. 22, 2356–2367 (2012).

36. Hou, F. et al. MAVS forms functional prion-like aggregates to activate and propagate antiviral innate immune response. Cell 146, 448–461 (2011).

37. Schüler, A. & Bornberg-Bauer, E. Evolution of Protein Domain Repeats in Metazoa. Mol. Biol. Evol. 33, 3170–3182 (2016).

38. Zeng, W. et al. Reconstitution of the RIG-I Pathway Reveals a Signaling Role of Unanchored Polyubiquitin Chains in Innate Immunity. Cell 141, 315–330 (2010).

39. Gack, M. U. et al. TRIM25 RING-finger E3 ubiquitin ligase is essential for RIG-I-mediated antiviral activity. Nature 446, 916–920 (2007).

40. Kouwaki, T., Nishimura, T., Wang, G., Nakagawa, R. & Oshiumi, H. K63_-_linked polyubiquitination of LGP2 by Riplet regulates RIG_-_I_-_dependent innate immune response. EMBO Rep. 24, e54844 (2023).

41. Peisley, A., Wu, B., Xu, H., Chen, Z. J. & Hur, S. Structural basis for ubiquitin-mediated antiviral signal activation by RIG-I. Nature 509, 110–114 (2014).

42. Wu, B. et al. Molecular imprinting as a signal-activation mechanism of the viral RNA sensor RIG-I. Mol. Cell 55, 511–523 (2014).

43. Wu, B. & Hur, S. How RIG-I like receptors activate MAVS. Curr. Opin. Virol. 12, 91–98 (2015).

44. Fitzgerald, M. E., Rawling, D. C., Vela, A. & Pyle, A. M. An evolving arsenal: viral RNA detection by RIG-I-like receptors. Curr. Opin. Microbiol. 20, 76–81 (2014).

45. Onomoto, K., Onoguchi, K. & Yoneyama, M. Regulation of RIG-I-like receptor-mediated signaling: interaction between host and viral factors. Cell. Mol. Immunol. 18, 539–555 (2021).

46. Pett, W. et al. The Role of Homology and Orthology in the Phylogenomic Analysis of Metazoan Gene Content. Mol. Biol. Evol. 36, 643–649 (2019).

47. Nestor, B. J., Bayer, P. E., Fernandez, C. G. T., Edwards, D. & Finnegan, P. M. Approaches to increase the validity of gene family identification using manual homology search tools. Genetica 151, 325–338 (2023).

48. Bernheim, A., Cury, J. & Poirier, E. Z. The immune modules conserved across the tree of life: Towards a definition of ancestral immunity. PLoS Biol. 22, e3002717 (2024).

49. Wein, T. & Sorek, R. Bacterial origins of human cell-autonomous innate immune mechanisms. Nat. Rev. Immunol. 22, 629–638 (2022).

50. Zipple, M. N., Vogt, C. C. & Sheehan, M. J. Re-wilding Model Organisms: Opportunities to test causal mechanisms in social determinants of health and aging. Neurosci. Biobehav. Rev. 152, 105238 (2023).

51. Oyesola, O. et al. Genetic and environmental interactions contribute to immune variation in rewilded mice. Nat. Immunol. 25, 1270–1282 (2024).

52. Barber, G. N. Host defense, viruses and apoptosis. Cell Death Differ. 8, 113–126 (2001).

53. Orzalli, M. H. & Kagan, J. C. Apoptosis and necroptosis as host defense strategies to prevent viral infection. Trends Cell Biol. 27, 800–809 (2017).

54. Wein, T. et al. CARD domains mediate anti-phage defence in bacterial gasdermin systems. Nature 639, 727–734 (2025).

55. Genikhovich, G. & Technau, U. Induction of spawning in the starlet sea anemone Nematostella vectensis, in vitro fertilization of gametes, and dejellying of zygotes. Cold Spring Harb. Protoc. 2009, pdb.prot5281 (2009).

56. Hand, C. & Uhlinger, K. R. The Culture, Sexual and Asexual Reproduction, and Growth of the Sea Anemone Nematostella vectensis. Biol. Bull. 182, 169–176 (1992).

57. Ikmi, A., McKinney, S. A., Delventhal, K. M. & Gibson, M. C. TALEN and CRISPR/Cas9-mediated genome editing in the early-branching metazoan Nematostella vectensis. Nat. Commun. 5, 5486 (2014).

58. Concordet, J.-P. & Haeussler, M. CRISPOR: intuitive guide selection for CRISPR/Cas9 genome editing experiments and screens. Nucleic Acids Res. 46, W242–W245 (2018).

59. Zhorov, B. S. Possible Mechanism of Ion Selectivity in Eukaryotic Voltage-Gated Sodium Channels. J. Phys. Chem. B 125, 2074–2088 (2021).

60. Karabulut, A., He, S., Chen, C.-Y., McKinney, S. A. & Gibson, M. C. Electroporation of short hairpin RNAs for rapid and efficient gene knockdown in the starlet sea anemone, Nematostella vectensis. Dev. Biol. 448, 7–15 (2019).

61. Admoni, Y., Kozlovski, I., Lewandowska, M. & Moran, Y. TATA Binding Protein (TBP) Promoter Drives Ubiquitous Expression of Marker Transgene in the Adult Sea Anemone Nematostella vectensis. Genes 11, 1081 (2020).

62. Schindelin, J., et al. Fiji: an open-source platform for biological-image analysis. Nat. Methods 9, 676–682 (2012).

63. Aguilar-Camacho, J. M. et al. Functional analysis in a model sea anemone reveals phylogenetic complexity and a role in cnidocyte discharge of DEG/ENaC ion channels. Commun. Biol. 6, 17 (2023).

64. Wang, H., He, M., Willard, B. & Wu, Q. Cross-linking, Immunoprecipitation and Proteomic Analysis to Identify Interacting Proteins in Cultured Cells. Bio-Protoc. 9, e3258 (2019).

65. Data | Carnegie Coral & Marine Organisms. https://cmo.carnegiescience.edu/data.

66. Acropora muricata genome assembly ASM3666990v1. NCBI https://www.ncbi.nlm.nih.gov/datasets/genome/GCF_036669905.1/.

67. aiptasiav2@reefgenomics: genomes of three Aiptasia strains. http://aiptasiav2.reefgenomics.org/.

68. Zimmermann, B. et al. Sea anemone genomes reveal ancestral metazoan chromosomal macrosynteny. Preprint at 10.1101/2020.10.30.359448 (2020).

69. Tamura, K., Stecher, G. & Kumar, S. MEGA11: Molecular Evolutionary Genetics Analysis Version 11. Mol. Biol. Evol. 38, 3022–3027 (2021).

70. Nguyen, L.-T., Schmidt, H. A., von Haeseler, A. & Minh, B. Q. IQ-TREE: A Fast and Effective Stochastic Algorithm for Estimating Maximum-Likelihood Phylogenies. Mol. Biol. Evol. 32, 268–274 (2015).

71. Guindon, S. et al. New Algorithms and Methods to Estimate Maximum-Likelihood Phylogenies: Assessing the Performance of PhyML 3.0. Syst. Biol. 59, 307–321 (2010).

72. Bolger, A. M., Lohse, M. & Usadel, B. Trimmomatic: a flexible trimmer for Illumina sequence data. Bioinforma. Oxf. Engl. 30, 2114–2120 (2014).

73. Putnam, N. H. et al. Sea anemone genome reveals ancestral eumetazoan gene repertoire and genomic organization. Science 317, 86–94 (2007).

74. Dobin, A. et al. STAR: ultrafast universal RNA-seq aligner. Bioinforma. Oxf. Engl. 29, 15–21 (2013).

75. Liao, Y., Smyth, G. K. & Shi, W. featureCounts: an efficient general purpose program for assigning sequence reads to genomic features. Bioinforma. Oxf. Engl. 30, 923–930 (2014).

76. Love, M. I., Huber, W. & Anders, S. Moderated estimation of fold change and dispersion for RNA-seq data with DESeq2. Genome Biol. 15, 550 (2014).

77. Robinson, M. D., McCarthy, D. J. & Smyth, G. K. edgeR: a Bioconductor package for differential expression analysis of digital gene expression data. Bioinforma. Oxf. Engl. 26, 139–140 (2010).

78. Chen, H. & Boutros, P. C. VennDiagram: a package for the generation of highly-customizable Venn and Euler diagrams in R. BMC Bioinformatics 12, 35 (2011).

79. Wu, T. et al. clusterProfiler 4.0: A universal enrichment tool for interpreting omics data. Innov. Camb. Mass 2, 100141 (2021).

80. Binns, D. et al. QuickGO: a web-based tool for Gene Ontology searching. Bioinforma. Oxf. Engl. 25, 3045–3046 (2009).

81. Frickey, T. & Lupas, A. CLANS: a Java application for visualizing protein families based on pairwise similarity. Bioinforma. Oxf. Engl. 20, 3702–3704 (2004).

82. Alboukadel, K. & Fabian, M. factoextra: Extract and Visualize the Results of Multivariate Data Analyses. CRAN Contrib. Packag. 10.32614/cran.package.factoextra(2016) doi:10.32614/cran.package.factoextra.

83. Katoh, K. & Standley, D. M. MAFFT multiple sequence alignment software version 7: improvements in performance and usability. Mol. Biol. Evol. 30, 772–780 (2013).

84. Minh, B. Q. et al. IQ-TREE 2: New Models and Efficient Methods for Phylogenetic Inference in the Genomic Era. Mol. Biol. Evol. 37, 1530–1534 (2020).

85. Katoh, K. & Standley, D. M. MAFFT Multiple Sequence Alignment Software Version 7: Improvements in Performance and Usability. Mol. Biol. Evol. 30, 772–780 (2013).

86. Yang, Z. PAML 4: Phylogenetic Analysis by Maximum Likelihood. Mol. Biol. Evol. 24, 1586–1591 (2007).

87. Mirdita, M. et al. ColabFold: making protein folding accessible to all. Nat. Methods 19, 679–682 (2022).

88. Babraham Bioinformatics - FastQC A Quality Control tool for High Throughput Sequence Data. https://www.bioinformatics.babraham.ac.uk/projects/fastqc/.

89. The genome sequence of the starlet sea … | Wellcome Open Research. https://wellcomeopenresearch.org/articles/8-79.

90. Kim, D., Paggi, J. M., Park, C., Bennett, C. & Salzberg, S. L. Graph-based genome alignment and genotyping with HISAT2 and HISAT-genotype. Nat. Biotechnol. 37, 907–915 (2019).

91. Danecek, P. et al. Twelve years of SAMtools and BCFtools. GigaScience 10, giab008 (2021).

92. Langmead, B., Trapnell, C., Pop, M. & Salzberg, S. L. Ultrafast and memory-efficient alignment of short DNA sequences to the human genome. Genome Biol. 10, R25 (2009).

93. Quast, C. et al. The SILVA ribosomal RNA gene database project: improved data processing and web-based tools. Nucleic Acids Res. 41, D590–596 (2013).

94. Nawrocki, E. P. et al. Rfam 12.0: updates to the RNA families database. Nucleic Acids Res. 43, D130–137 (2015).

95. Parks, D. H. et al. GTDB: an ongoing census of bacterial and archaeal diversity through a phylogenetically consistent, rank normalized and complete genome-based taxonomy. Nucleic Acids Res. 50, D785–D794 (2022).

96. Bushmanova, E., Antipov, D., Lapidus, A. & Prjibelski, A. D. rnaSPAdes: a de novo transcriptome assembler and its application to RNA-Seq data. GigaScience 8, giz100 (2019).

