## Supplementary Figures for "An ancient anthozoan protein reveals an alternative evolutionary path of antiviral signaling"

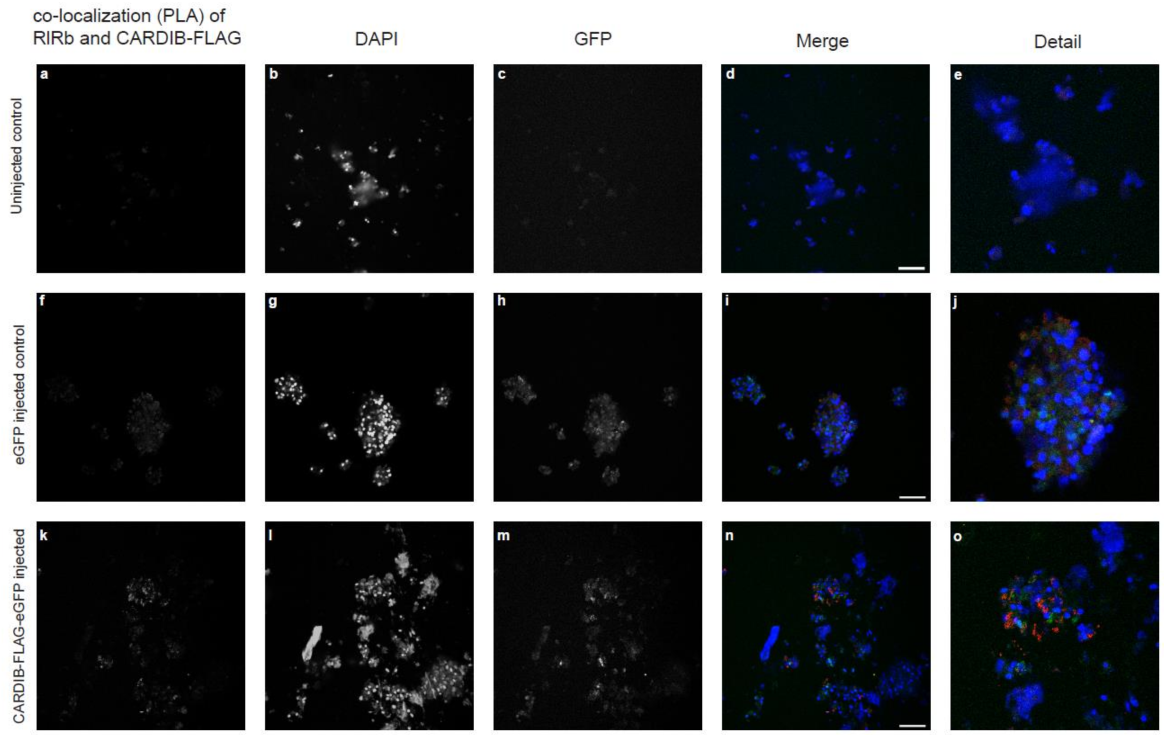


**Supplementary Fig 1.** Confocal images of PLA signals in dissociated cells derived from uninjected 24-hour-old planulae and injected with eGFP mRNA control and C-terminal CARDIB-FLAG-GFP mRNA, respectively; PLA signal for RLRb and FLAG colocalization (red); DAPI (blue); GFP (green); scale bars 20 μm. Representative single z-stack section images were obtained from at least three independent replicates. **a-e,** cells derived from uninjected planulae, used as the control group. **f-j,** cells derived from planulae that were injected with eGFP mRNA, used as the control group. **k-o,** cells derived from planulae that were injected with C-terminal CARDIB-FLAG-GFP mRNA.

N-terminal FLAGx3-CARDIB-eGFP Nve gBlock

T7 promoter ef1aKozak CARDIB P2A mCherry FLAG GFP complexity fix 3'UTR

TAATACGACTCACTATAAGTGTTAAACCAACCAACCACC**ATG**GGCGGGTCAGACTACAAAGATGACGA
TGATAAGGGAGGCTCCGACTATAAGGATGACGATGACAAAGGTGGAAGCGATTACAAGGACGATGATG
ATAAAGGGGGCAGTGATAGAGTTCTGCCAGACATTGTGGAAGATGTTATACCTGGAGACATGATGCCA
TACTTACCCTGCCTAACCGACGATGACAAAGAGCAAATTTTGTGTGAGGAGGAGAACAGGGGGAGCAG
AAGAGCAGCTTATCTACTTGTTGATAGACTCAAACGGAGAAGGAATGGCATGTTTGACTTTATTAGGG
CATTGAGTAAAACAGGCTGTCACCATGTTGTAGCTCGTATTGATGAGGAAATCCAAAAACAGAATTAC
AGACAACCGCAACCCGGAGGCTCAGGCGCTACGAATTTCAGTCTGCTGAAGCAAGCGGGCGACGTGGA
AGAGAATCCAGGCCCTGGAGGTTCGGTGAGCAAGGGCGAGGAGCTGTTCACCGGGGTGGTGCCCATCC
TGGTCGAGCTGGACGGCGACGTAAACGGCCACAAGTTCAGCGTGTCCGGCGAGGGCGAGGGCGATGCC
ACCTACGGCAAGCTGACCCTGAAGTTCATCTGCACCACCGGCAAGCTGCCCGTGCCCTGGCCCACCCT
CGTGACCACCCTGACCTACGGCGTGCAGTGCTTCAGCCGCTACCCCGACCACATGAAGCAGCACGACT
TCTTCAAGTCCGCCATGCCCGAAGGCTACGTCCAGGAGCGCACCATCTTCTTCAAGGACGACGGCAAC
TACAAGACCCGCGCCGAGGTGAAGTTCGAGGGCGACACCCTGGTGAACCGCATCGAGCTGAAGGGCAT
CGACTTCAAGGAGGACGGCAACATCCTGGGGCACAAGCTGGAGTACAACTACAACAGCCACAACGTCT

ATATCATGGCCGACAAGCAGAAGAACGGCATCAAGGTGAACTTCAAGATCCGCCACAACATCGAGGAC
GGCAGCGTGCAGCTCGCCGACCACTACCAGCAGAACACCCCCATCGGCGACGGCCCCGTGCTGCTGCC
CGACAACCACTACCTGAGCACCCAGTCCGCCCTGAGCAAAGACCCCAACGAGAAGCGCGATCACATGG
TCCTGCTGGAGTTCGTGACCGCCGCCGGGATCACTCTCGGCATGGACGAGCTGTACAAG**TAA**GACTCT
AGATCATAATCAGCCATACCACATTTGTAGAGGTTTTACTTGCTTTAAAAAACCTGAAACATACCCAC
ACCTCCCCCTGAACCTGAAACATAAAATGAATGCAATTGTTGTTGGAAACATATTAACTTGTTTATTG
CAGCTTATAATGGTTACAAATGAAAGCAATAGCATCACAAATTTCACAAATAGAAGCATTTTTTTCAC
TGC

C-terminal CARDIB-FLAGx3-eGFP Nve gBlock

TAATACGACTCACTATAAGTGTTAAACCAACCAACCACC**ATG**GATAGAGTTCTGCCAGACATTGTGGA
AGATGTTATACCTGGAGACATGATGCCATACTTACCCTGCCTAACCGACGATGACAAAGAGCAAATTT
TGTGTGAGGAGGAGAACAGGGGGAGCAGAAGAGCAGCTTATCTACTTGTTGATAGACTCAAACGGAGA
AGGAATGGCATGTTTGACTTTATTAGGGCATTGAGTAAAACAGGCTGTCACCATGTTGTAGCTCGTAT
TGATGAGGAAATCCAAAAACAGAATTACAGACAACCGCAACCCGGCGGGTCAGACTACAAAGATGACG
ATGATAAGGGAGGCTCCGACTATAAGGATGACGATGACAAAGGTGGAAGCGATTACAAGGACGATGAT
GATAAAGGGGGCAGTGGAGGCTCAGGCGCTACGAATTTCAGTCTGCTGAAGCAAGCGGGCGACGTGGA
AGAGAATCCAGGCCCTGGAGGTTCGGTGAGCAAGGGCGAGGAGCTGTTCACCGGGGTGGTGCCCATCC
TGGTCGAGCTGGACGGCGACGTAAACGGCCACAAGTTCAGCGTGTCCGGCGAGGGCGAGGGCGATGCC
ACCTACGGCAAGCTGACCCTGAAGTTCATCTGCACCACCGGCAAGCTGCCCGTGCCCTGGCCCACCCT
CGTGACCACCCTGACCTACGGCGTGCAGTGCTTCAGCCGCTACCCCGACCACATGAAGCAGCACGACT
TCTTCAAGTCCGCCATGCCCGAAGGCTACGTCCAGGAGCGCACCATCTTCTTCAAGGACGACGGCAAC
TACAAGACCCGCGCCGAGGTGAAGTTCGAGGGCGACACCCTGGTGAACCGCATCGAGCTGAAGGGCAT
CGACTTCAAGGAGGACGGCAACATCCTGGGGCACAAGCTGGAGTACAACTACAACAGCCACAACGTCT
ATATCATGGCCGACAAGCAGAAGAACGGCATCAAGGTGAACTTCAAGATCCGCCACAACATCGAGGAC
GGCAGCGTGCAGCTCGCCGACCACTACCAGCAGAACACCCCCATCGGCGACGGCCCCGTGCTGCTGCC
CGACAACCACTACCTGAGCACCCAGTCCGCCCTGAGCAAAGACCCCAACGAGAAGCGCGATCACATGG
TCCTGCTGGAGTTCGTGACCGCCGCCGGGATCACTCTCGGCATGGACGAGCTGTACAAG**TAA**GACTCT
AGATCATAATCAGCCATACCACATTTGTAGAGGTTTTACTTGCTTTAAAAAACCTGAAACATACCCAC
ACCTCCCCCTGAACCTGAAACATAAAATGAATGCAATTGTTGTTGGAAACATATTAACTTGTTTATTG
CAGCTTATAATGGTTACAAATGAAAGCAATAGCATCACAAATTTCACAAATAGAAGCATTTTTTTCAC
TGC

eGFP gBlock

TAATACGACTCACTATAAGTGTTAAACCAACCAACCACC**ATG**GTGAGCAAGGGCGAGGAGCTGTTCAC
CGGGGTGGTGCCCATCCTGGTCGAGCTGGACGGCGACGTAAACGGCCACAAGTTCAGCGTGTCCGGCG
AGGGCGAGGGCGATGCCACCTACGGCAAGCTGACCCTGAAGTTCATCTGCACCACCGGCAAGCTGCCC
GTGCCCTGGCCCACCCTCGTGACCACCCTGACCTACGGCGTGCAGTGCTTCAGCCGCTACCCCGACCA
CATGAAGCAGCACGACTTCTTCAAGTCCGCCATGCCCGAAGGCTACGTCCAGGAGCGCACCATCTTCT
TCAAGGACGACGGCAACTACAAGACCCGCGCCGAGGTGAAGTTCGAGGGCGACACCCTGGTGAACCGC
ATCGAGCTGAAGGGCATCGACTTCAAGGAGGACGGCAACATCCTGGGGCACAAGCTGGAGTACAACTA

CAACAGCCACAACGTCTATATCATGGCCGACAAGCAGAAGAACGGCATCAAGGTGAACTTCAAGATCC
GCCACAACATCGAGGACGGCAGCGTGCAGCTCGCCGACCACTACCAGCAGAACACCCCCATCGGCGAC
GGCCCCGTGCTGCTGCCCGACAACCACTACCTGAGCACCCAGTCCGCCCTGAGCAAAGACCCCAACGA
GAAGCGCGATCACATGGTCCTGCTGGAGTTCGTGACCGCCGCCGGGATCACTCTCGGCATGGACGAGC
TGTACAAG**TAA**GACTCTAGATCATAATCAGCCATACCACATTTGTAGAGGTTTTACTTGCTTTAAAAA
ACCTGAAACATACCCACACCTCCCCCTGAACCTGAAACATAAAATGAATGCAATTGTTGTTGGAAACA
TATTAACTTGTTTATTGCAGCTTATAATGGTTACAAATGAAAGCAATAGCATCACAAATTTCACAAAT
AGAAGCATTTTTTTCACTGC

**Supplementary Fig 2.** The sequence and the design of the N-terminal FLAGx3-CARDIB-eGFP Nve gBlock.

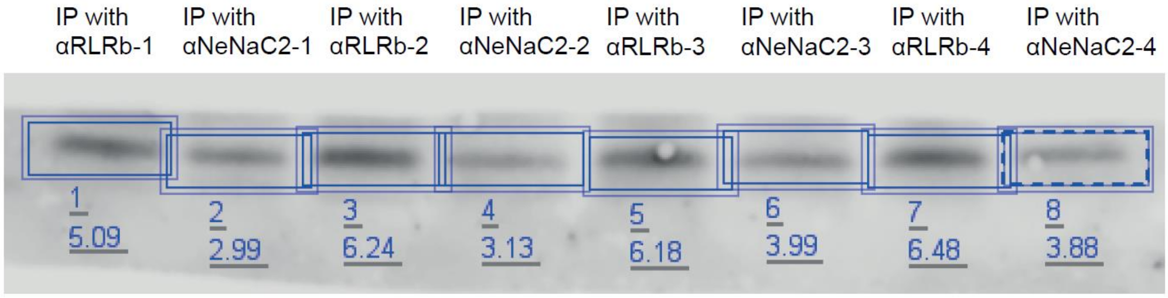


**Supplementary Fig 3.** Western blot image showing CARDIB presence, after Co-Immunoprecipitation of RLRb with CARDIB from *N. vectensis* 24-h embryos as described in the materials and methods section. Membrane was incubated with αCARDIB antibody as primary antibody and Peroxidase conjugated antibody against rabbit as secondary antibody and finally detected with a CCD camera of the Odyssey Fc imaging system (Li-COR Biosciences) using Clarity™ ECL and Clarity™ max kits (Biorad) as described in detail. Co-IP Sample name and biological repeat number are indicated above the blot image. The upper numbers below the blot image show the serial numbers of the bands on the blot. The lower numbers below the blot image indicate band intensities as were measured by the Image Studio software (Li-COR Biosciences) and used for the fold change calculations between RLRb and NeNaC2 samples in each repeat.

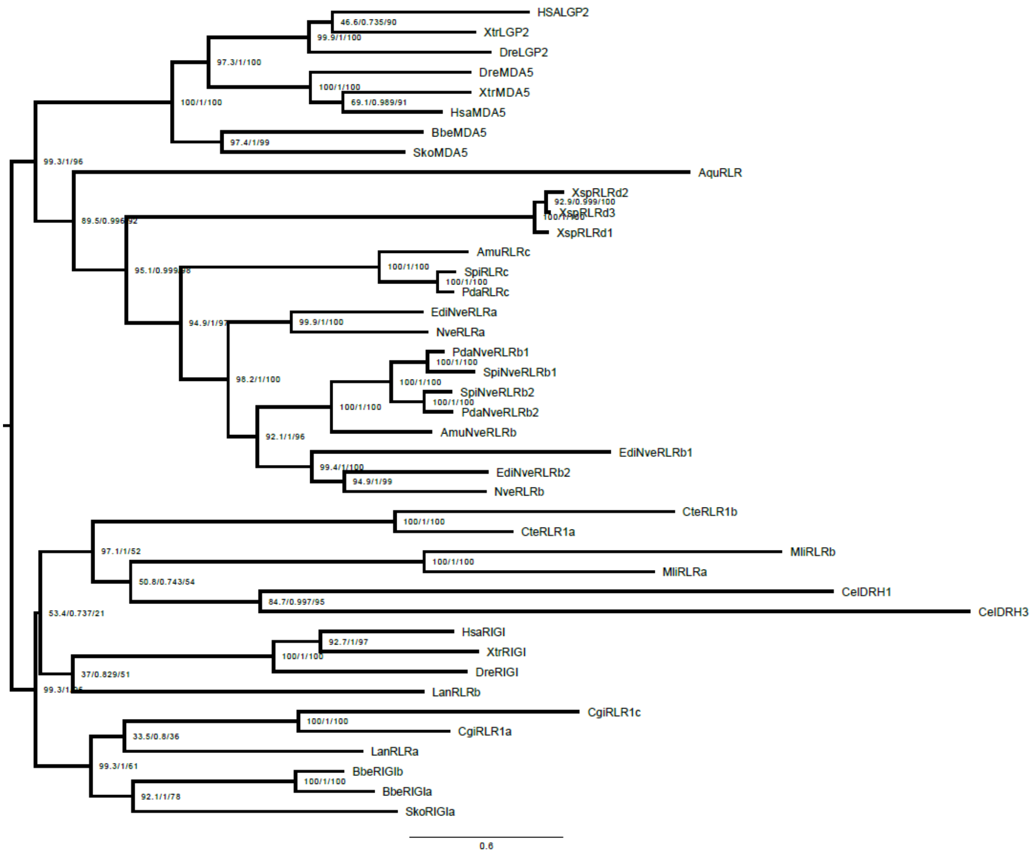


**Supplementary Fig 4.** Phylogenetic relationship of metazoan RLRs. Maximum likelihood consensus phylogenetic tree of representative RLR sequences, numbers in parentheses are SH-aLRT support (%) / aBayes support / ultrafast bootstrap support (%). Amu, *Acropora muricata;* Aqu, *Amphimedon queenslandica*; Bbe, *Branchiostoma belcheri*; Cte, *Capitella teleta*; Cgi, *Crassostrea gigas*; Cel, *Caenorhabditis elegans*; Dre, *Danio rerio*; Edia, *Exaiptasia diaphana*; Hsa, *Homo sapiens*; Lan, *Lingula anatina*; Mli, *Macrostomum lignano*; Nve, *Nematostella vectensis*; Pda, *Pocillopora damicronis*; Spi, *Stylophora pistillata*; Xsp, *Xenia spp; Xtr*, *Xenopus tropicalis*.

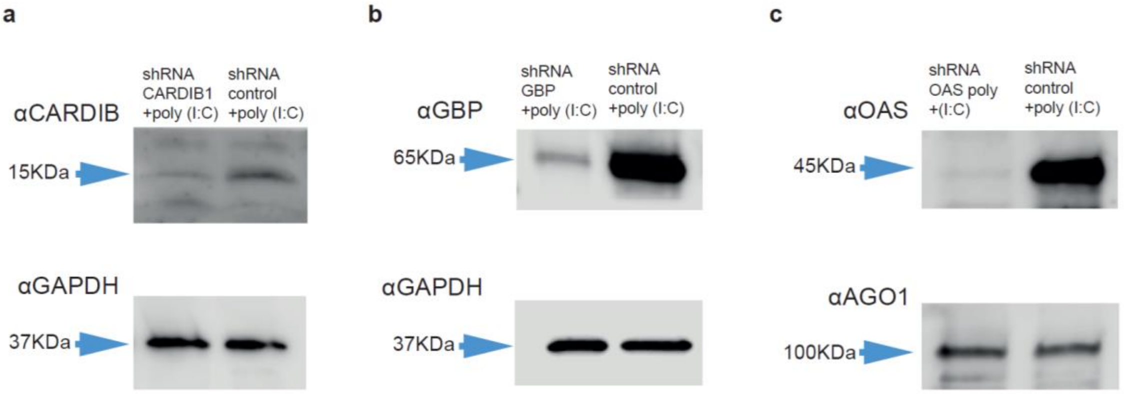


**Supplementary Fig 5.** Antibodies validation of CARDIB, GBP, and OAS via western blot. Zygotes were injected with shRNA against each of the target genes in addition to poly (I:C), and the gene expression was tested at the protein level. Scramble shRNA with poly (I:C) injected as a control. **a,** extracted protein from planulae that were injected with shRNA with poly (I:C) against CARDIB and scramble shRNA with poly (I:C) as a control. **b,** extracted protein from planulae that were injected with shRNA with poly (I:C) against GBP and scramble shRNA with poly (I:C) as a control. **c,** extracted protein from planulae that were injected with shRNA with poly (I:C) against OAS and scramble shRNA with poly (I:C) as a control. For the normalization of the amount of protein in the western blot, GAPDH was used for the samples of αCARDIB and αGBP, for the samples of αOAS, AGO1 was used for the normalization of the amount of protein to avoid size similarity (which was shown not to respond to poly (I:C) or be involved in the *N. vectensis* immunity[1]) used for the normalization.

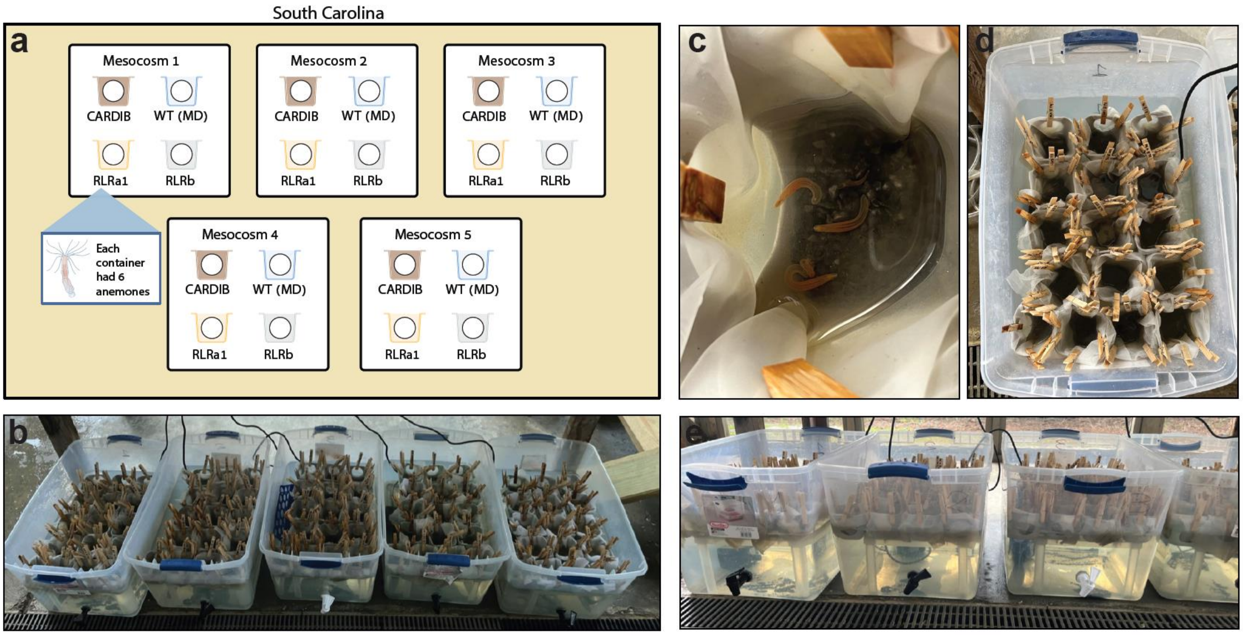


**Supplementary Fig 6.** Mesocosm Field Design, **a,** Cartoon depiction of the mesocosm samples used in this study (created with BioRender.com). Each Mesocosm (plastic bin) is a replicate. Animal container placements were randomized in each replicate. **b**, The five mesocosm replicates in the field at Belle W. Baruch Marine Field Laboratory (Georgetown, South Carolina, US). **c,** A close-up image of the inside of an animal container. Each animal container contained one *N. vectensis* strain and began with six individuals to be sampled at different time points. **d**, An above view of the inside of one mesocosm. **e**, A side view of the mesocosms to demonstrate the water line. Natural estuary water was added to submerge half of the animal containers to prevent animals from escaping.

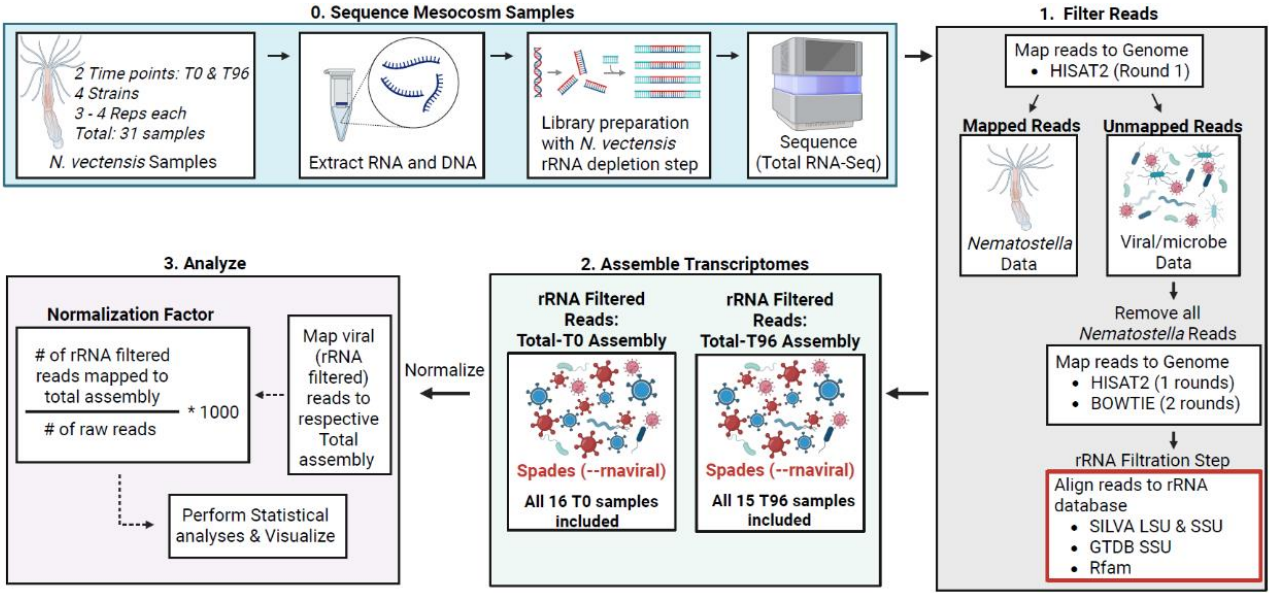


**Supplementary Fig 7. Bioinformatic pipeline to assess viral read counts.** *N. vectensis* samples were collected at two timepoints during the mesocosm experiment: the initial timepoint (T0), prior to exposure, and after 96 hours of exposure to natural estuary water (T96). Four strains were examined with 3-4 biological replicates each, yielding a total of 31 samples (i.e., four replicates for each T0 and T96 strain, except for CARDIB T96, which had three replicates). RNA was extracted from each sample and used to generate RNA libraries for total RNA sequencing.

In the bioinformatics pipeline, the first step after adapter trimming and quality control (FastQC) was host read removal. Trimmed reads were iteratively aligned to the *N. vectensis* genome until no reads mapped, leaving viral and microbial reads. An rRNA filtration step was then performed by aligning these unmapped reads to an rRNA database to remove prokaryotic and eukaryotic rRNA sequences. The resulting rRNA-filtered dataset was designated as “presumably viral reads.”

These reads were used to assemble transcriptomes. Two assemblies were generated: the Total-T0 assembly, constructed from all 16 T0 samples, and the Total-T96 assembly, constructed from all 15 T96 samples. Viral reads from each sample were then mapped back to the appropriate assembly, and normalization factors were calculated. Statistical analyses were subsequently performed. Figure schematic created with BioRender.com.
